# Strain-Dependent Kinetic Properties of KIF3A and KIF3C Tune the Mechanochemistry of the KIF3AC Heterodimer

**DOI:** 10.1101/793075

**Authors:** Brandon M. Bensel, Michael S. Woody, Serapion Pyrpassopoulos, Yale E. Goldman, Susan P. Gilbert, E. Michael Ostap

## Abstract

KIF3AC is a mammalian neuron-specific kinesin-2 implicated in intracellular cargo transport. It is a heterodimer of KIF3A and KIF3C motor polypeptides which have distinct biochemical and motile properties as engineered homodimers. Single-molecule motility assays show that KIF3AC moves processively along microtubules at a rate faster than expected given the motility rates of the KIF3AA and much slower KIF3CC homodimers. To resolve the stepping kinetics of KIF3A and KIF3C motors in homo-and heterodimeric constructs, and to determine their transport potential under mechanical load, we assayed motor activity using interferometric scattering (iSCAT) microscopy and optical trapping. The distribution of stepping durations of KIF3AC molecules is described by a rate (*k*_*1*_ = 11 s^−1^) without apparent kinetic asymmetry in stepping. Asymmetry was also not apparent under hindering or assisting mechanical loads of 1 pN in the optical trap. KIF3AC shows increased force sensitivity relative to KIF3AA, yet is more capable of stepping against mechanical load than KIF3CC. Microtubule gliding assays containing 1:1 mixtures of KIF3AA and KIF3CC result in speeds similar to KIF3AC, indicating the homodimers mechanically impact each other’s motility to reproduce the behavior of the heterodimer. We conclude that the stepping of KIF3C can be activated by KIF3A in a strain-dependent manner which is similar to application of an assisting load, and the behavior of KIF3C mirrors prior studies of kinesins with increased interhead compliance. These results suggest that KIF3AC-based cargo transport likely requires multiple motors, and its mechanochemical properties arise due to the strain-dependences of KIF3A and KIF3C.

**Significance Statement:** Kinesins are important long-range intracellular transporters in neurons required by the extended length of the axon and dendrites and selective cargo transport to each. The mammalian kinesin-2, KIF3AC, is a neuronal heterodimer of fast and slow motor polypeptides. Our results show that KIF3AC has a single observed stepping rate in the presence and absence of load and detaches from the microtubule rapidly under load. Interestingly, both KIF3A and assisting loads accelerate the kinetics of KIF3C. These results suggest that KIF3AC is an unconventional cargo transporter and its motile properties do not represent a combination of alternating fast and slow step kinetics. We demonstrate that the motile properties of KIF3AC represent a mechanochemistry that is specific to KIF3AC and may provide functional advantages in neurons.

## Introduction

Kinesin-2 is a distinctive subfamily of processive kinesins that contains both homodimeric and heterodimeric motors involved in microtubule (MT) plus-end directed cargo transport (1–4). The mammalian heterodimeric kinesin-2s result from three genes: *kif3a, kif3b,* and *kif3c* to form heterodimeric motors KIF3AB and KIF3AC (5–9) while expression of the *kif17* gene product results in homodimeric kinesin-2, KIF17 (10–12). Moreover, KIF3A and KIF3B do not form homodimers, and KIF3B does not heterodimerize with KIF3C suggesting distinct transport functions of KIF3AB, KIF3AC, and KIF17. While these studies show that KIF3AB and KIF3AC heterodimerization is preferred over homodimerization, there is evidence for an injury-specific homodimer of KIF3CC in neurons (13). KIF3C also contains a signature 25-residue insert of glycines and serines in loop L11 of the catalytic motor domain, which has been shown to regulate KIF3AC processivity (14) and microtubule catastrophe promoted by KIF3CC *in vitro* (15). The structural diversity within the kinesin-2 subfamily suggests that heterodimerization of kinesin-2 motors may be a mechanism to optimize the mechanochemistry of the motors for specific tasks (16).

Motility and biochemical experiments indicate that the activities of KIF3A and KIF3C depend on the motors with which they are partnered. KIF3AC and the engineered homodimers, KIF3AA and KIF3CC, move processively in single-molecule motility assays. KIF3AC has longer run lengths than either of the homodimers (14), and its speed is substantially faster than expected given the very slow rate of the KIF3CC homodimer (see Discussion). Additionally, comparison of the presteady-state kinetics of KIF3AC, KIF3AA, and KIFCC indicates that the rates of association of the motors with microtubules and ADP release depend on the motor’s partner head. These kinetic and motile properties suggest that interhead communication within the heterodimer tunes the mechanochemical properties of KIF3A and KIF3C (4, 14, 17, 18).

We sought to resolve the stepping kinetics of individual KIF3A and KIF3C motor domains in homo-and heterodimeric motors and to determine the transport potential of the dimers under mechanical load. To accomplish this, we assayed motor activity using interferometric scattering (iSCAT) microscopy, a recent advance in light microscopy which has sufficient spatiotemporal precision to resolve single steps of a kinesin motor during processive runs (19–23), and optical trapping which reveals the motor performance under forces hindering and assisting plus-end directed motility (24–29). We observed that the step dwell time distribution of KIF3AC is not a combination of the dwell time distributions observed for KIF3AA and KIF3CC and is adequately fit by a single exponential rate. The symmetric stepping of KIF3AC persists under hindering or assisting mechanical load. We show that the kinetics of KIF3C are accelerated by assisting load, and an equal mixture of KIF3AA and KIF3CC promotes microtubule gliding at the same rate as KIF3AC. Therefore, our results suggest that the strain-dependent kinetics of KIF3A and KIF3C give rise to the velocity of KIF3AC. KIF3AC shows decreased stall and detachment forces relative to KIF3AA and detaches from the microtubule rapidly under hindering or assisting mechanical load. We argue that the mechanochemical properties of KIF3AC observed here and previously are adaptations for cargo transport in large ensembles, and that single or even a few KIF3AC motors are unlikely to transport cargo against sustained load in a similar manner to kinesin-1.

## Results

### KIF3AA, KIF3AC, and KIF3CC step dwell times and backstepping frequencies are distinct in unloaded conditions

We resolved the stepping displacements and kinetics of KIF3AA, KIF3CC, and KIF3AC using interferometric scattering (iSCAT) microscopy. Streptavidin-coated gold nanoparticles (50 nm) were attached to the C-terminal His-tags of kinesin dimers via a biotinylated anti-His antibody to measure the movement of single motors as the alternating heads interacted with a surface-attached microtubule (*Supplemental Appendix,* Fig. S1*A*).

In the presence of 1 mM MgATP, KIF3AA moved processively at a rate of 270 ± 75 nm/s (Fig. 1*A and B*), which is similar to the velocity measured previously with the same protein construct in single molecule Qdot motility assays (14), but slightly slower than observed in optical trapping assays that used a construct with a different dimerization motif (16). Individual steps were clearly resolved in the iSCAT traces (Fig. 1*B*). After filtering by a Chung-Kennedy filter (30) with a 20 ms width, traces were fit using a step-finding algorithm ((31); Fig. 1*B*) to extract the distribution of dwell times and step sizes (Fig. 2*A and B*). A double Gaussian distribution was fit to the forward step sizes, and single Gaussian distribution was fit to the backward steps. The most prominent component is centered near 8 nm (9.1 ± 3.5 nm; Fig. 2*B* and Table 1) with a smaller population (16 ± 15 nm) likely representing two steps that occurred rapidly in sequence and could not be resolved by the step-fitting algorithm. Infrequent backward steps were also resolved (−14 ± 7.2 nm; A_back_ = 0.12). The dwell times, plotted as cumulative distribution functions (CDFs) (Fig. 2*A*), were best fit by the sum of two exponential functions (Fig. 2*A* and Table 1). The fit revealed a predominant fast component (A_1_ = 0.83) with a rate of *k*_1_ = 34 s^−1^. The slower, minor component, *k*_2_ = 6.1 s^−1^ (A_2_ = 0.17), is most likely due to infrequent pauses in stepping (Fig. 1*B*, asterisks).

**Table 1.** iSCAT step kinetics and displacements fit parameters. The relative amplitudes and rates of each exponential fit are shown. Single exponential fit equation: 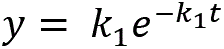 and sum of two exponential functions fit equation: 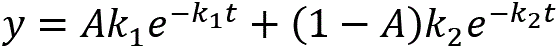 where the amplitude A is reported as A_1_ and (1 − A) is reported as A_2_. For step displacements, A_back_ and μ_back_ describe backstepping, A_forward1_ and μ_forward1_ the dominant forward step size peak near 8 nm, and A_forward2_ and μ_forward2_ the additional forward peak near 16 nm. Backward and forward step size distributions were fit independently. For the KIF3AA dwell time fit, the log-likelihood test gives p = 1.4 × 10^−7^ justifying a double exponential fit, and for the KIF3CC dwell time fit, the log-likelihood ratio test gives p = 0.036. Means are shown ± σ of the fit curve where σ indicates the width of the Gaussian peak and not the uncertainty in peak position. Dashes, not applicable.

| Motor | [MgATP] | Step Kinetics |  |  |  | Step Sizes |  |  |  |  |  |
| --- | --- | --- | --- | --- | --- | --- | --- | --- | --- | --- | --- |
|  |  | A<sub>1</sub> | k<sub>1</sub> (s<sup>-1</sup>) | A<sub>2</sub> | k<sub>2</sub> (s<sup>-1</sup>) | A<sub>back</sub> | μ<sub>back</sub> (nm) | A<sub>forward1</sub> | μ<sub>forward1</sub> (nm) | A<sub>forward2</sub> | μ<sub>forward2</sub> (nm) |
| KIF3AA | 1 mM | 0.83 | 34 | 0.17 | 6.1 | 0.12 | -14 ± 7.2 | 0.49 | 9.1 ± 3.5 | 0.39 | 16 ± 15 |
| KIF3CC | 1 mM | 0.68 | 1.3 | 0.32 | 0.44 | 0.18 | -8.2 ± 7.2 | 0.66 | 6.9 ± 2.9 | 0.16 | 19 ± 8.2 |
| KIF3AC | 1 mM | 1.0 | 11 | — | — | 0.15 | -12 ± 8.2 | 0.60 | 9.3 ± 3.4 | 0.25 | 16 ± 6.4 |

**Fig. 1:**
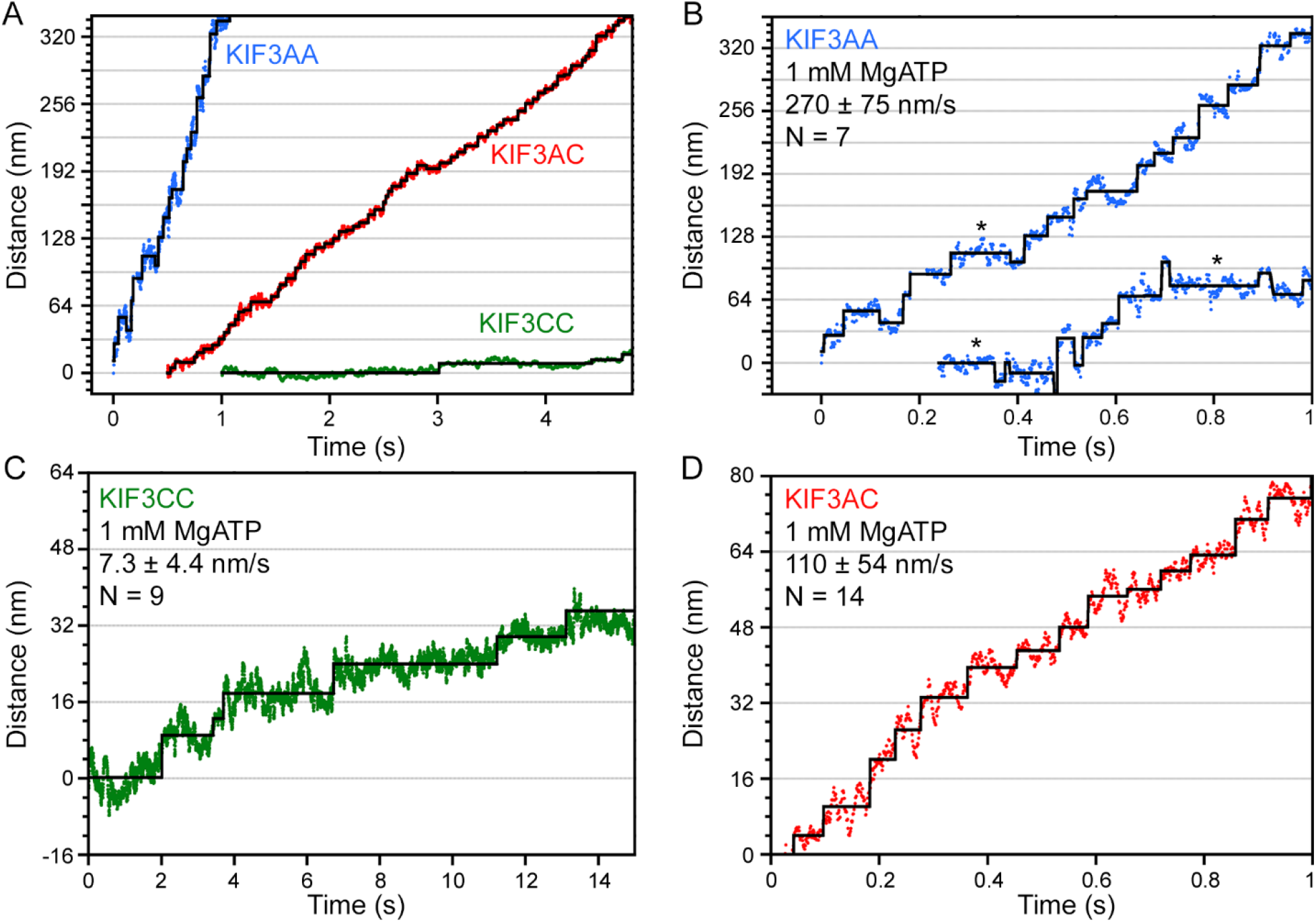
Nanometric tracking of KIF3AC and engineered homodimers by iSCAT microscopy. (A) Sample traces of KIF3AA (blue), KIF3AC (red), and KIF3CC (green) tracked by iSCAT microscopy shown on the same time and length scales for comparison. Sample traces (B-D) of expanded views of the position along path of travel at 1 mM MgATP for KIF3AA (B), KIF3CC (C), and KIF3AC (D). An asterisk (*) is used to denote instances of pausing for >150 ms in traces of KIF3AA motility. Velocity is reported as the mean ± S.D. for each motor. N values reflect the number of molecules tracked for iSCAT analysis. Black overlay, Kerssemakers’ algorithm fit (31).

**Fig. 2:**
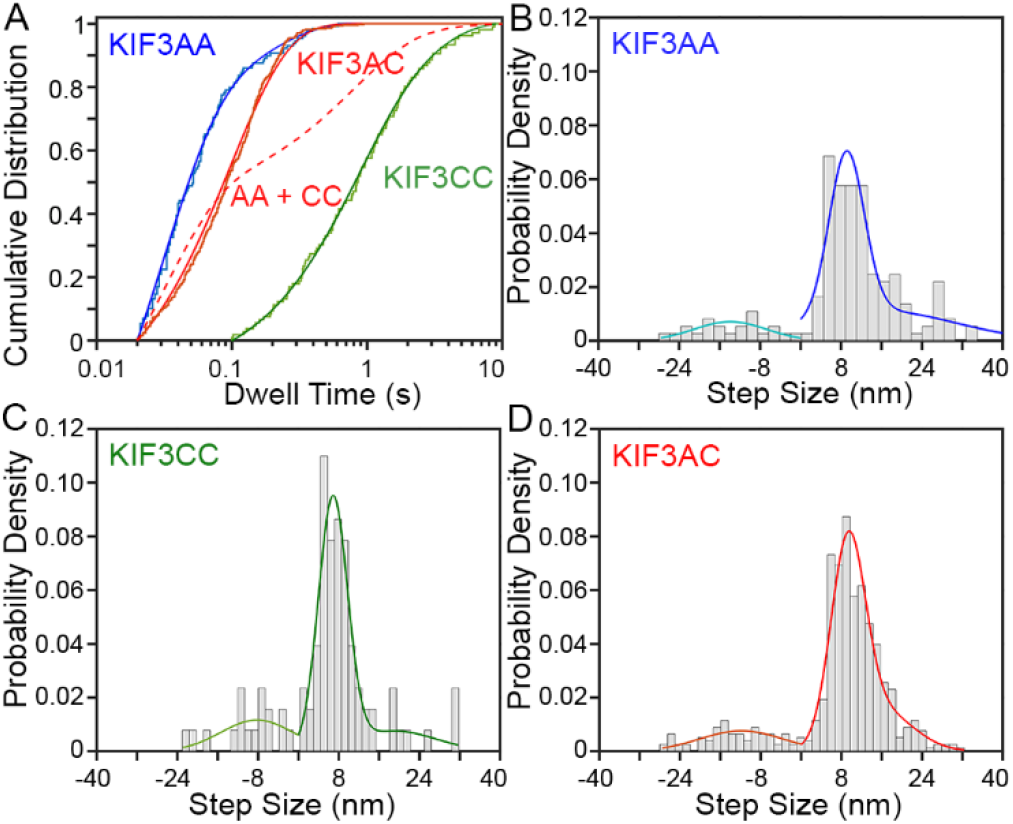
KIF3AC steps by a single rate at 1 mM MgATP with 8-nm steps. Step sizes and kinetics measured by iSCAT microscopy for KIF3AC and engineered homodimers. (A) Cumulative distribution function plots and accompanying fits of dwell time data at 1 mM MgATP for KIF3AA (blue), KIF3AC (red), and KIF3CC (green). Dwell time data are fit to a single exponential function (KIF3AA) or the sum of two exponential functions as justified by the log-likelihood ratio test (KIF3CC, KIF3AC). Hypothetical curve of the sum of the KIF3AA and KIF3CC fits is plotted (red dashed line). Data are plotted with a logarithmic x-axis. (B-D) Histograms of step sizes observed for KIF3AA (B), KIF3CC (C), and KIF3AC (D). Negative step sizes refer to backward steps. Forward and backward step size distributions were fit independently, and fits are shown in different colors. Fit parameters are given in Table 1. For KIF3AA N_steps_ = 168, for KIF3CC N_steps_ = 93, and for KIF3AC N_steps_ = 516.

KIF3CC homodimers moved much slower than KIF3AA under identical conditions with an average velocity (7.3 ± 4.4 nm/s, Fig. 1*A and C*) similar to values measured previously (14). Step-fitting to the KIF3CC traces revealed a step size distribution (Fig. 2*C* and Table 1) with a prominent component (A_forward1_ = 0.66) centered at 6.9 ± 2.9 nm. Backward steps (−8.2 ± 7.2 nm; A_back_ = 0.18; Table 1) occurred more frequently than observed for KIF3AA as isolated steps and not backward runs. There were fewer steps detected near 16 nm, consistent with fewer unresolved fast steps. The distribution of observed dwell times for KIF3CC was best fit by the sum of two exponential functions (green, Fig. 2*A*) with rates *k*_*1*_= 1.3 s^−1^ and *k*_*2*_ = 0.44 s^−1^ (A_1_ = 0.68, A_2_ = 0.32; Table 1). The ~30-fold difference in stepping rates between KIF3AA and KIF3CC is consistent with previous single-molecule velocity reports (4, 14).

KIF3AC heterodimers moved processively at a velocity of 110 ± 54 nm/s (Fig. 1*D*). As observed for KIF3AA and KIF3CC, the most prominent distribution of step sizes for KIF3AC was centered near 8 nm (9.3 ± 3.4 nm; Fig. 2*D*, Table 1). Backsteps (−12 ± 8.2 nm, A_back_ = 0.15) were also observed for KIF3AC, as single isolated steps and not processive backward runs. Notably, the cumulative distribution of dwell times for KIF3AC (red, Fig. 2*A*) was best fit by a single-exponential function (*k*_*1*_ = 11 s^−1^; (32)) rather than the sum of two exponentials, which may have been expected given the ~30-fold difference in the unloaded stepping rates of KIF3A and KIF3C homodimers. Mathematical modeling was used to determine the conditions under which a single exponential fit could result from two independent rates. We found that at least a 2-fold difference in rates was necessary to statistically justify fitting the sum of two exponential components (*Supplemental Appendix* Fig. S4). The difference between observed and predicted dwell time distributions is illustrated by plotting a hypothetical CDF that is the normalized sum of KIF3AA and KIF3CC dwell times (Fig. 2*A*, red dashed curve). The data are clearly different from such a model.

### KIF3AC detaches from microtubules at lower forces than kinesin-2 KIF3AB or kinesin-1 KIF5B

We used a stationary, single-beam, optical trap to determine the ability of heterodimeric KIF3AC and the engineered KIF3AA and KIF3CC homodimers to move and remain attached to the microtubule in the presence of hindering loads. The unitary detachment, maximum, and stall forces of KIF3AC, KIF3AA, and KIF3CC bound to 0.82-μm streptavidin-coated beads via a biotinylated anti-His antibody were measured (see *Supplemental Appendix,* **SI Methods,** Fig. S1*B* and *C*). We also determined the same parameters for KIF3AB using this experimental geometry to compare with other published results (16, 33, 34).

The detachment and stall forces of KIF3AA (3.1 – 3.3 pN) are slightly lower than those of KIF3AB (Fig. 3*A-D* and Table 2), which are similar to values reported previously (16, 33, 34). KIF3CC and KIF3AC have substantially lower detachment-forces than KIF3AA (Fig. 3*C-H* and Table 2) at F_detach_ = 1.5 ± 0.4 pN for KIF3CC and F_detach_ = 1.9 ± 0.5 pN for KIF3AC. The maximum force and stall force measured for KIF3AA are greater than those for KIF3AC which are in turn greater than those for KIF3CC. A small percentage of events terminate in a stall (10-14%) for KIF3AB, KIF3AA, and KIF3AC, and no stall plateaus were observed for KIF3CC (Table 2). KIF3 motors are more likely to detach under hindering load rather than to become stalled. Previous work showed that a unique insert in loop L11 of KIF3C plays a role in regulating the run length of KIF3AC such that KIF3AC with the KIF3C loop L11 truncated to the length observed in KIF3B (KIF3ACΔL11) was more processive than native KIF3AC (14). However, no significant differences in the detachment, maximum, and stall forces or stall percentage between KIF3AC and KIF3ACΔL11 were observed (Table 2 and *Supplemental Appendix* Fig. S2). This result suggests that force sensitivity, as measured in this experimental geometry, is not modulated by loop L11.

**Table 2.** Measured ramp force parameters for KIF3 motors. Table of detachment, maximum, and stall force and stall percentage for KIF3AB, KIF3AA, KIF3CC, KIF3AC, and KIF3ACΔL11. Displayed values are means ± S.D. Maximum force is defined as the largest force value over baseline that occurs between the beginning of the force ramp and the detachment event. The stall force is defined as the mean force over the final 70 ms of an event that comes to a stall, where a stall is defined as a force plateau in which the standard deviation of the force is less than or equal to 5% of the mean force over that window (34). Dashes represent parameters that were not observed. Measured parameters for KIF3AC and KIF3ACΔL11 were compared through an unpaired student’s T-test with an α-reliability level of 0.05, and the differences between KIF3AC and KIF3ACΔL11 for F_detach_, F_max_, and F_stall_ were not significant (P=1, 0.36, 0.10 respectively). Measured detachment forces of KIF3AB, KIF3AA, KIF3AC and KIF3CC were compared through an unpaired Student’s t-test with an α-reliability level of 0.05, and all differences were highly significant (P<0.0001). Stall % was compared for KIF3AC and KIF3ACΔL11 using the “N-1” Chi-squared test, and no significant difference was observed (P=0.81).

| | $F_{\text{Detachment}}$<br>(pN) | $F_{\text{Maximum}}$<br>(pN) | $F_{\text{Stall}}$<br>(pN) | Stall % |
| --- | --- | --- | --- | --- |
| KIF3AB | $3.7 \pm 1.4$ | $4.2 \pm 1.4$ | $5.2 \pm 0.2$ | 10 |
| KIF3AA | $3.1 \pm 1.0$ | $3.3 \pm 1.1$ | $3.8 \pm 0.1$ | 13 |
| KIF3CC | $1.5 \pm 0.4$ | $1.6 \pm 0.4$ | — | — |
| KIF3AC | $1.9 \pm 0.5$ | $2.3 \pm 0.6$ | $2.6 \pm 0.5$ | 14 |
| KIF3ACΔL11 | $1.9 \pm 0.7$ | $2.4 \pm 1.2$ | $2.3 \pm 0.3$ | 13 |

**Fig. 3:**
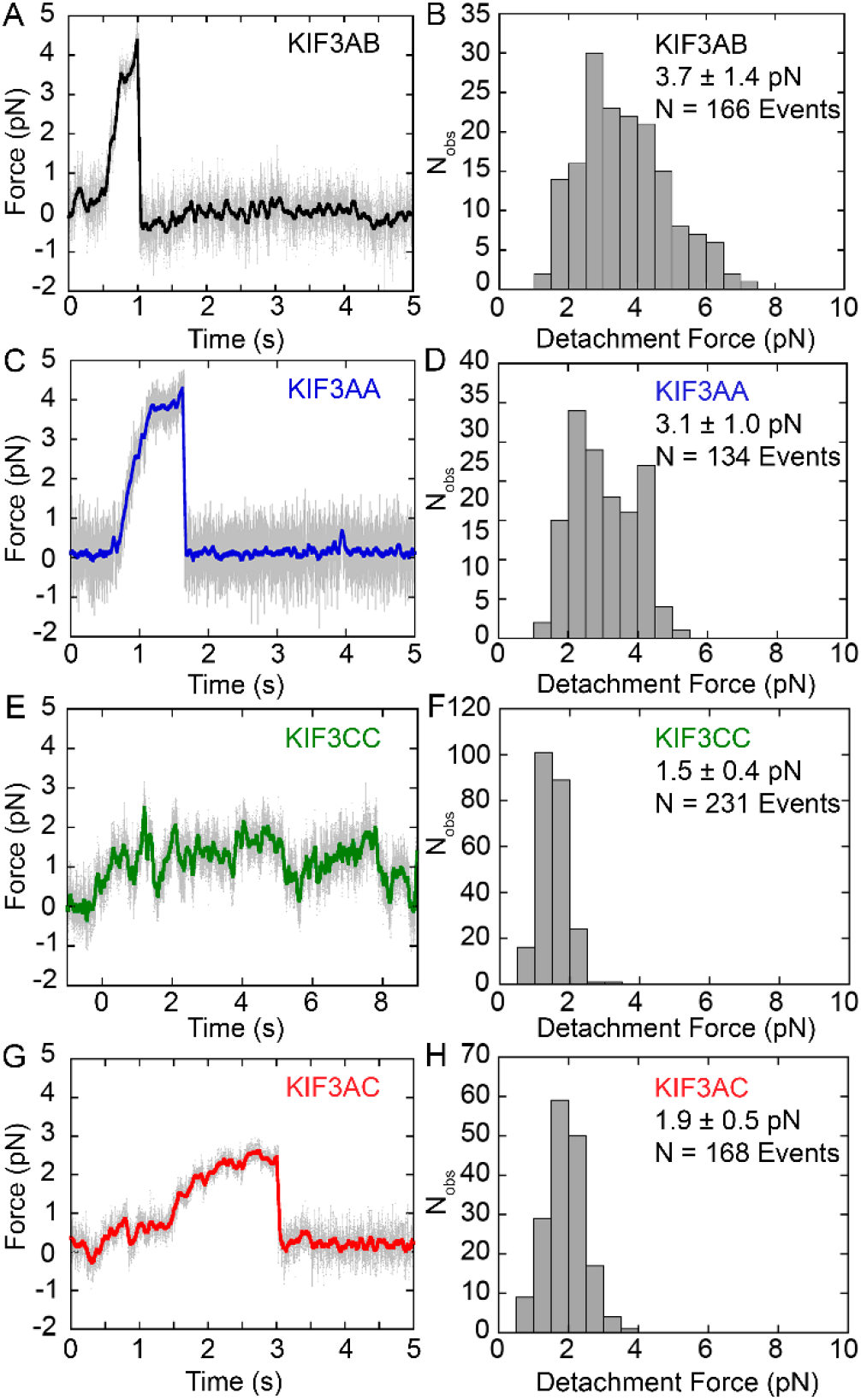
Force traces and detachment forces measured for KIF3 motor dimers. (A, C, E, G) Sample force ramps produced by each KIF3 dimer stepping against a stationary trap. Force increases as the distance between the bead center and trap center increases. Raw data are plotted in grey and colored overlaid data are filtered with a 20 ms mean filter for visualization. (B, D, F, H) Respective histograms of detachment forces. Detachment force is defined as the amount of force on the bead immediately prior to detachment, shown as mean ± S.D. N values reflect number of force ramps analyzed. (A, B) KIF3AB, (C, D) KIF3AA, (E, F) KIF3CC, and (G, H) KIF3AC.

### KIF3AC motility parameters are highly sensitive to hindering and assisting external loads

We determined the effect of constant hindering or assisting loads on the velocity, run length, and stepping kinetics of KIF3AA, KIF3CC, and KIF3AC using force-feedback in the optical trap (Figs. 4-5 and Table 3). KIF3AA stepped at an average velocity of 140 ± 64 nm/s under 1-pN hindering load and 190 ± 85 nm/s under 1-pN assisting load (Fig. 4*A*, 5*A* and Table 3). KIF3AA motility was observed over the range of 2-pN assisting loads to 4-pN hindering external loads, and the change in velocity with force was approximately 2-fold over this range. This force sensitivity is reflected by the effective distance parameter derived from the fit of the Bell equation 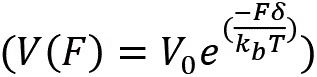 (35) to the data, δ = 0.59 ± 0.05 nm (Fig. 5*A*). The effective distance parameter of KIF3AA represents a relatively weak force dependence for velocity compared to conventional kinesin-1, δ = 3.58 nm, as was reported previously (16). Externally applied 1-pN assisting or hindering loads markedly decreased the average run length of KIF3AA relative to the run length in the absence of external load measured with Qdots ((14), Fig. 5*B* and Table 3). In contrast to KIF3AA, KIF3CC motility was nearly stalled under all hindering loads tested (0.5 to 2 pN, Fig. 4*B*, Fig. 5*C*, and Table 3). Assisting loads >1 pN resulted in faster motility by KIF3CC (Fig. 5*C* and Table 3). Much like KIF3AA, the KIF3CC run length was decreased >10-fold under both assisting and hindering loads compared to unloaded Qdot results ((14), unfilled square, Fig. 5*D*).

**Table 3.** Measured velocity and run length of KIF3 motors. Table of measured run lengths and velocities under 1-pN assisting load, unloaded, and 1-pN hindering load conditions at 1 mM MgATP. Displayed values are means ± S.D. Unloaded values are included from previous Qdot single molecule studies at 1 mM MgATP (14).

| Motor | 1 pN Assisting |  | Unloaded |  | 1 pN Hindering |  |
| --- | --- | --- | --- | --- | --- | --- |
|  | Vel (nm/s) | RL (nm) | Vel (nm/s) | RL (nm) | Vel (nm/s) | RL (nm) |
| KIF3AA | 190 ± 85 | 93 ± 68 | 240 ± 76 | 980 ± 910 | 140 ± 64 | 73 ± 62 |
| KIF3CC | 41 ± 22 | 54 ± 54 | 7.5 ± 6.3 | 570 ± 470 | 4.8 ± 16 | 25 ± 48 |
| KIF3AC | 170 ± 110 | 100 ± 57 | 170 ± 67 | 1300 ± 1700 | 68 ± 50 | 43 ± 48 |

**Fig. 4:**
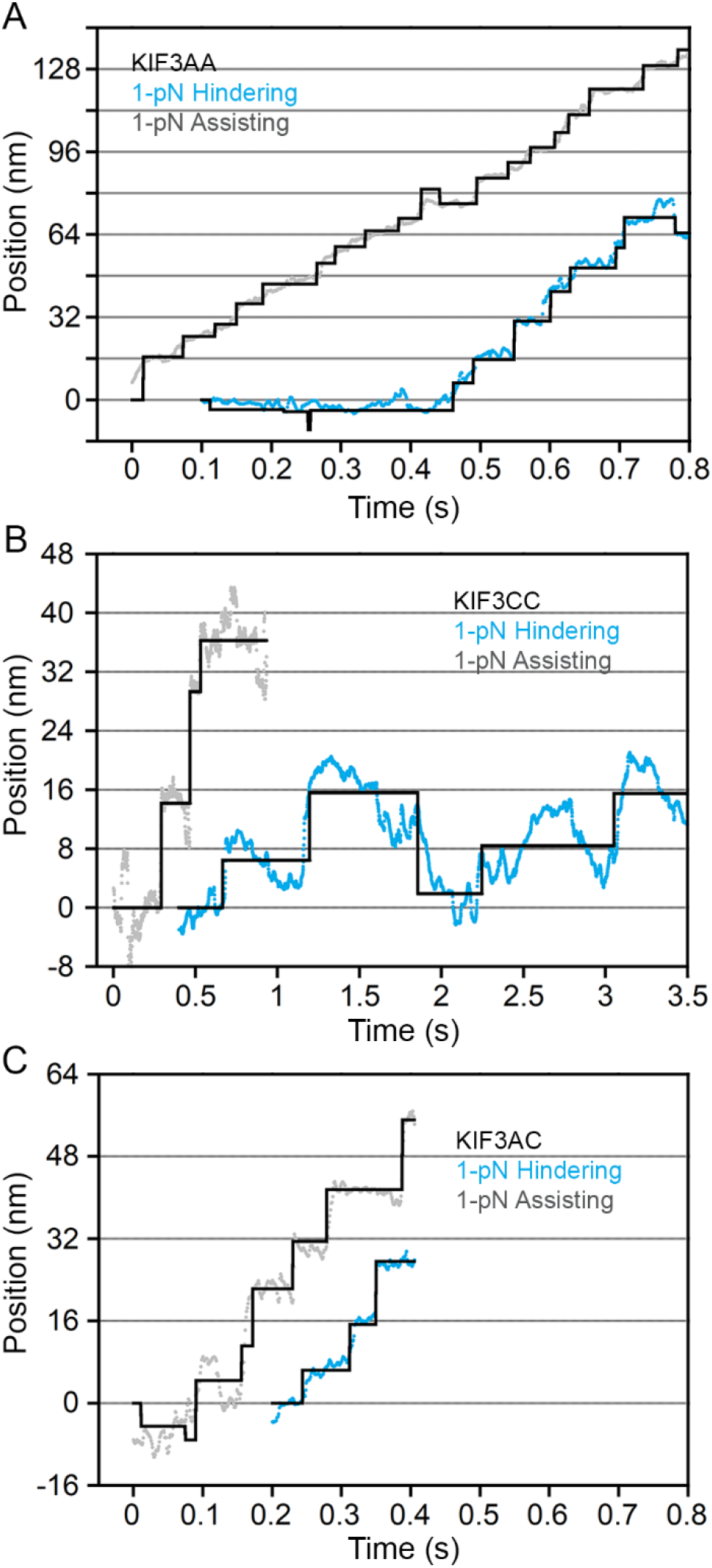
Stepping traces with constant assisting and hindering loads. Sample traces of the position of the optical trap as it maintained constant force on KIF3AA (A), KIF3CC (B), and KIF3AC (C) stepping against constant 1-pN hindering load (blue) and with constant 1-pN assisting load (grey). Black line: Kerssemakers’ algorithm fit to the data (31).

**Fig. 5:**
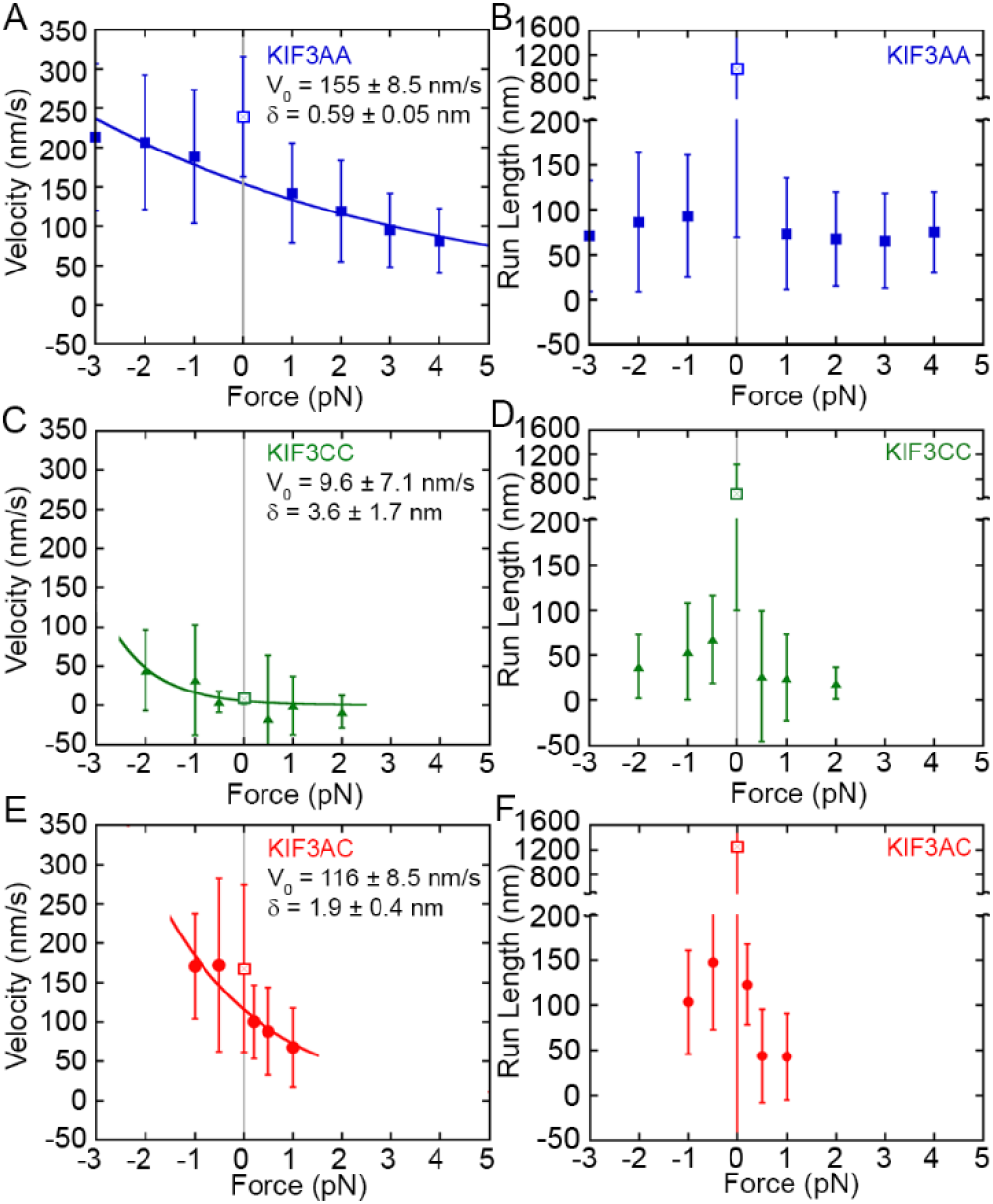
Force-dependence of velocity and run length measured for KIF3 motors. (A, C, E) Velocities ± S.D. versus load for KIF3AA (A), KIF3CC (C), and KIF3AC (E). Negative values on x-axis indicate assisting load, and positive values indicate hindering load. The Bell equation (35) fit to the trapping data is shown by a solid line on each plot: 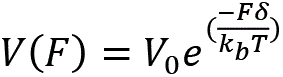 where *F* is the applied force in pN, *V* is the velocity, *V*_0_ is the velocity at zero load, *δ* is the effective distance parameter in nm, and k_B_T is the Boltzmann constant times the absolute temperature. A larger distance parameter indicates higher load sensitivity. Parameters in the absence of force, 0 on x-axis, are included from previous Qdot single molecule studies (14) and shown by an unfilled square with 0 on the x-axis highlighted by grey vertical lines. (B, D, F) Run lengths ± S.D. versus load for KIF3AA (B), KIF3CC (D), and KIF3AC (F). Negative values on the x-axis indicate assisting load, and positive values indicate hindering load.

The KIF3AC velocity load dependence (δ = 1.9 ± 0.4 nm; Fig. 5*E*) is larger than that of KIF3AA. Under a 1-pN hindering load, KIF3AC moved at a mean velocity of 68 ± 50 nm/s, yet application of a 1-pN assisting load resulted in an increased mean velocity of 170 ± 110 nm/s (Fig. 4*C*, Table 3). Similar to KIF3AA and KIF3CC, the run length of KIF3AC was dramatically altered by external loads and decreased ~10-fold relative to the run length determined in unloaded conditions of the Qdot single molecule motility assay ((14); Fig. 5*F*). Our results show that KIF3AC, KIF3AA, and KIF3CC show similar run length decrease under load (Fig. 5*F* and Table 3). In contrast, the force-dependence of velocity varies significantly among these three motors.

### KIF3AC, KIF3AA, and KIF3CC stepping kinetics and step size distributions are distinct under hindering or assisting loads

To determine how step sizes and step kinetics are affected by mechanical loads, we performed step-finding analysis on the traces of KIF3AA, KIF3CC, and KIF3AC motility under 1-pN constant hindering or assisting load at 1 mM MgATP (Fig. 4, black overlays). For KIF3AA under a 1-pN hindering load, very few backsteps (A_back_ = 0.04, Table 4) were observed. Slightly more occurred with a 1-pN assisting load (A_back_ = 0.12, Fig. 6*C and D*, Table 4). The distribution of dwell times observed for KIF3AA under 1-pN hindering load was fit by a single exponential function with *k*_*1*_ = 18 s^−1^ (Fig. 6*A*, blue and Table 4). With a 1-pN assisting load, the dwell time distribution of KIF3AA shown in Fig. 6*B* (blue) was fit by a single exponential function with *k*_*1*_ = 24 s^−1^ (Table 4). Given the average step size of KIF3AA near 8 nm, the rates derived from each dwell time distribution for KIF3AA are in agreement with the measured velocities.

**Table 4.** Optical trap step kinetics and step displacement fit parameters. Dwell time distributions were fit by either a single exponential function or the sum of two exponential functions to determine step kinetics. Single exponential fit equation: 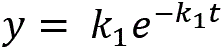 and sum of two exponential functions fit equation: 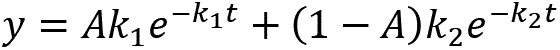 where the amplitude A is reported as A_1_ and (1 − A) is reported as A_2_. Amplitude and rate parameters are shown. Justification of a second exponential was determined by the log-likelihood ratio test. For KIF3CC, p < 1 × 10^−16^ at 1-pN hindering load, and p < 1 × 10^−16^ at 1-pN assisting load. For KIF3AC, p < 1 × 10^−16^ at 1-pN hindering load, and p < 1 × 10^−16^ at 1-pN assisting load. Step size distributions were fit by the sum of two Gaussian distributions. Relative amplitudes are reported; step sizes are reported as means ± σ of the fit curve, where σ indicates the width of Gaussian peak and not uncertainty in peak position.

| Motor | [MgATP] | Load (pN) | Step Kinetics |  |  |  | Step Sizes |  |  |  |
| --- | --- | --- | --- | --- | --- | --- | --- | --- | --- | --- |
|  |  |  | A<sub>1</sub> | k<sub>1</sub> (s<sup>-1</sup>) | A<sub>2</sub> | k<sub>2</sub> (s<sup>-1</sup>) | A<sub>back</sub> | μ<sub>back</sub> (nm) | A<sub>forward</sub> | μ<sub>forward</sub> (nm) |
| KIF3AA | 1 mM | 1 pN Hindering | 1.0 | 18 | — | — | 0.04 | -7.1 ± 4.8 | 0.96 | 8.7 ± 2.9 |
| KIF3AA | 1 mM | 1 pN Assisting | 1.0 | 24 | — | — | 0.12 | -11 ± 6.4 | 0.88 | 11 ± 5.5 |
| KIF3CC | 1 mM | 1 pN Hindering | 0.61 | 6.6 | 0.39 | 0.6 | 0.39 | -6.7 ± 5.2 | 0.61 | 7.8 ± 4.4 |
| KIF3CC | 1 mM | 1 pN Assisting | 0.78 | 18 | 0.22 | 2.2 | 0.20 | -9.8 ± 5.1 | 0.80 | 11 ± 7.5 |
| KIF3AC | 10 μM | 1 pN Hindering | 1.0 | 5.9 | — | — | 0.35 | -5.0 ± 7.2 | 0.65 | 7.8 ± 2.9 |
| KIF3AC | 1 mM | 1 pN Hindering | 0.96 | 11 | 0.04 | 2.4 | 0.21 | -7.4 ± 5.0 | 0.79 | 9.2 ± 4.5 |
| KIF3AC | 1 mM | 1 pN Assisting | 0.88 | 24 | 0.12 | 2.7 | 0.27 | -9.4 ± 7.4 | 0.73 | 11 ± 7.3 |

**Fig. 6:**
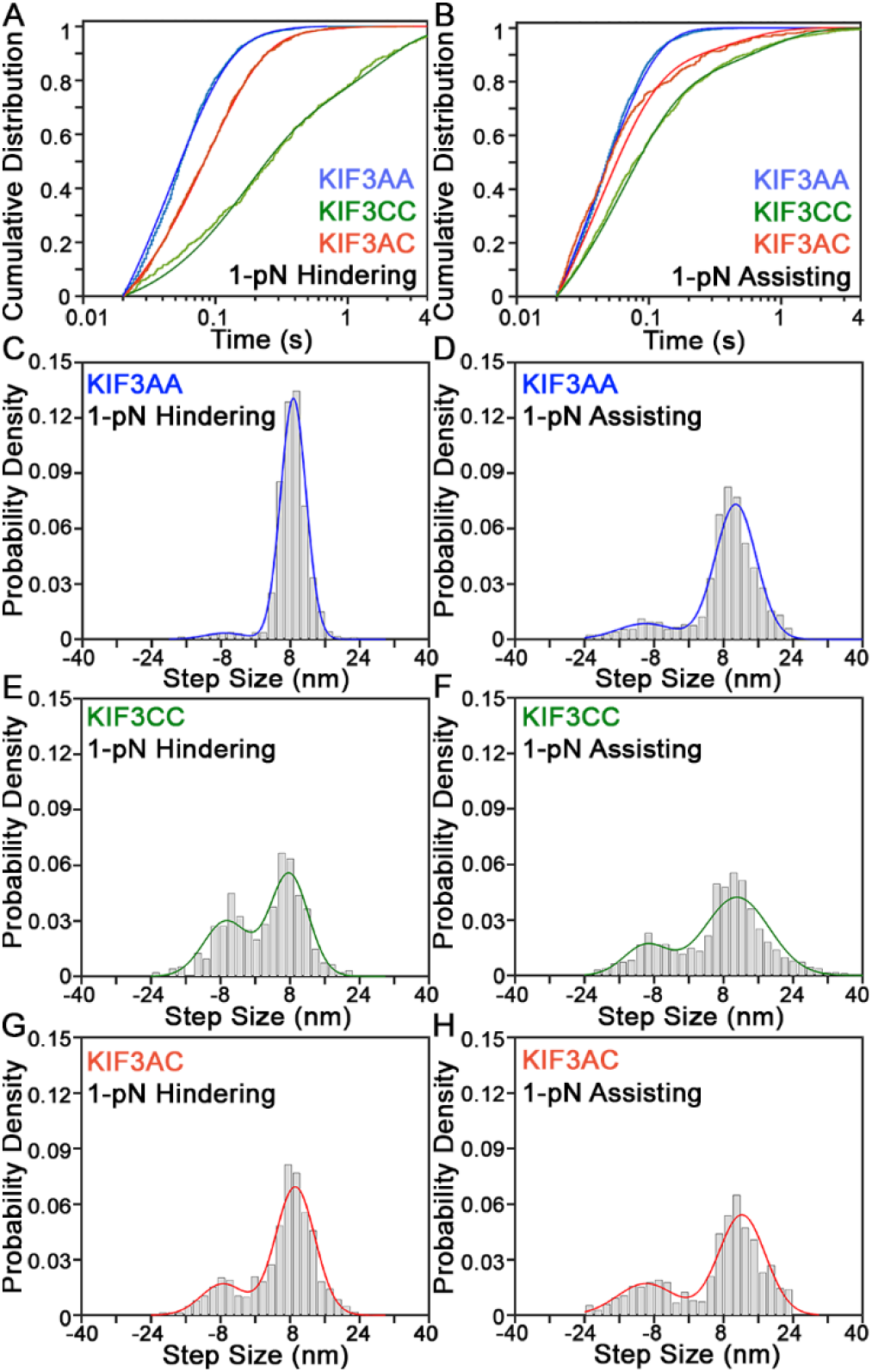
Step sizes and dwell time distributions derived from constant force stepping traces. (A) Cumulative distribution function plot of dwell times for KIF3AA (blue, N_steps_ = 1783), KIF3CC (green, N_steps_ = 502), and KIF3AC (red, N_steps_ = 1271) measured under a 1-pN hindering load. Lighter colored data are plotted with a darker fit line. (B) CDF plot of dwell times for KIF3AA (blue, N_steps_ = 2820), KIF3CC (green, N_steps_ = 1480), and KIF3AC (red, N_steps_ = 845) measured under a 1-pN assisting load. Dwell time distributions are fitted to a single exponential function (KIF3AA) or the sum of two exponential functions (KIF3CC, KIF3AC) as justified by the log-likelihood ratio test. Step size histograms are shown for 1-pN hindering or assisting load for KIF3AA (C, D), KIF3CC (E, F), and KIF3AC (G, H). Negative step sizes indicate backsteps toward the microtubule minus end, and positive step sizes indicate forward steps toward the microtubule plus end. Step size distributions are fit by the sum of two Gaussian distributions.

In contrast to KIF3AA, KIF3CC at 1-pN hindering load exhibited many backward steps. The step size distribution had peaks centered at both 7.8 ± 4.4 nm (A_forward_ = 0.61) and −6.7 ± 5.2 nm (A_back_ = 0.39) (Fig. 6*E and F*, Table 4). The proportion of backsteps taken by KIF3CC under hindering load (A_back_ = 0.39) was greater than that observed under assisting load (A_back_ = 0.20) or in the absence of load (A_back_ = 0.18, Table 1). The sum of two exponential functions provided the best fit to the cumulative distribution of dwell times observed for KIF3CC under 1-pN hindering and assisting load (Fig. 6*A and B*). Under hindering load, the KIF3CC dwell time distribution showed a fast rate *k*_*1*_ = 6.6 s^−1^ (A_1_ = 0.61), and a second slow rate, *k*_*2*_ = 0.6 s^−1^ (A_2_ = 0.39). In contrast, at 1-pN assisting load, these rates accelerated to *k*_*1*_= 18 s^−1^ (A_1_ = 0.78), and *k*_*2*_ = 2.2 s^−1^ (A_2_ = 0.22). In both cases, these rates were much slower than those observed for KIF3AA.

The step size distribution observed for KIF3AC at 1 mM MgATP (Fig. 6*G and H*) revealed a prominent component centered near 8 nm (9.2 ± 4.5 nm, A_forward_ = 0.79) under 1-pN hindering load and more abundant backsteps than observed in the absence of load (A_back_ = 0.21, Fig. 6*G and H*, Table 4). Step sizes observed for KIF3AC were similar under 1-pN assisting load and 1-pN hindering load. The cumulative distribution of dwell times for KIF3AC under 1-pN hindering and assisting loads are shown in red (Fig. 6*A and B*, respectively). Under a 1-pN hindering or assisting load, the dwell time distribution for KIF3AC was best fit by the sum of two exponential functions, based on the log-likelihood ratio test (32), with the majority of the dwell times associated with the faster rate. A rate of *k*_*1*_ = 11 s^−1^ (A_1_ = 0.96, Table 4; see *Supplemental Appendix* Results) described KIF3AC stepping under a 1-pN hindering load, and *k*_*1*_ = 24 s^−1^ (A_1_ = 0.88) described KIF3AC stepping with a 1-pN assisting load. As was observed in the absence of load, KIF3AA stepped with the fastest kinetics while KIF3CC stepped more slowly, and KIF3AC was intermediate to KIF3AA and KIF3CC.

KIF3AC motility at 10 μM MgATP under a 1-pN hindering load was also observed (*Supplemental Appendix* Fig. S3*A*). The step size distribution revealed primarily 8-nm forward steps (7.8 ± 2.9 nm, A_forward_= 0.65) with an additional population of backsteps (−5.0 ± 7.2 nm, A_back_ = 0.35) which was more prominent than that observed at 1 mM MgATP or in the absence of load (*Supplemental Appendix* Fig. S3*B*, Table 1 and Table 4). As was observed in zero load experiments, backsteps under load were typically single isolated step events. Surprisingly, the cumulative distribution of dwell times at 10 μM MgATP was adequately fit by a single exponential function with a rate constant *k_1_* = 5.9 s^−1^ (*Supplemental Appendix* Fig. S3*C*). KIF3AC did not show evidence for asymmetric stepping kinetics as would be expected from alternating fast and slow steps by the KIF3A and KIF3C heads. The relative frequency of backsteps varied between different motors at the MgATP concentrations tested.

### Mixed motor microtubule gliding assays recapitulate unloaded KIF3AC velocity

To determine if KIF3A and KIF3C motors can impact each other’s motility independently of heterodimerization, we performed microtubule gliding assays with mixtures of KIF3AA and KIF3CC. Assays were performed with combinations of motors that ranged in between 100% KIF3AA to 100% KIF3CC, keeping total motor concentration constant (Fig. 7*A*). KIF3AA alone propelled microtubules at a velocity of 210 ± 55 nm/s, and KIF3CC supported gliding at 1.9 ± 1.6 nm/s (Fig. 7*A* and *Supplemental Appendix* Fig. S5). Strikingly, the speed of microtubule gliding powered by an equal mixture of KIF3AA and KIF3CC was 120 ± 2.0 nm/s (Fig. 7*A* and *Supplemental Appendix* Fig. S5*C*), which is not significantly different from the gliding velocity of KIF3AC (126.4 ± 34.3 nm/s; P = 0.2037, unpaired student’s *t* test). An equal mixture of KIFAA and KIFCC in the gliding filament assay might simulate the internal strains in KIFAC to some extent. This is borne out by considering the strain dependent acceleration of KIFCC and the strain dependent slowing of KIFAA (Fig. 7*B*). At plus and minus 2.8 pN, the two homodimers are predicted to have equal velocities of 104 nm/s. Strikingly, the optical trap data for KIFAC intersect the zero-force axis at the very similar velocity of 110 nm/s (dashed curve in Fig. 7B). These results strongly indicate that mixtures of KIF3AA and KIF3CC homodimers mechanically impact each other’s motility in a way that reproduces the motile behavior of the KIF3AC heterodimer. The faster KIF3AA motor slows due to a hindering load imposed by the slower KIF3CC motor, while the velocity of KIF3CC increases due to the assisting load of KIF3AA. Strikingly, the relationship between microtubule gliding rate and motor ratio can be modeled directly (Fig. 7*A*) using the force-dependent parameters obtained from optical trapping results (Fig. 5; see Methods). Thus, the intramolecular strain and the force dependence of the two motors likely explain the motile behavior of KIF3AC heterodimer.

**Fig. 7:**
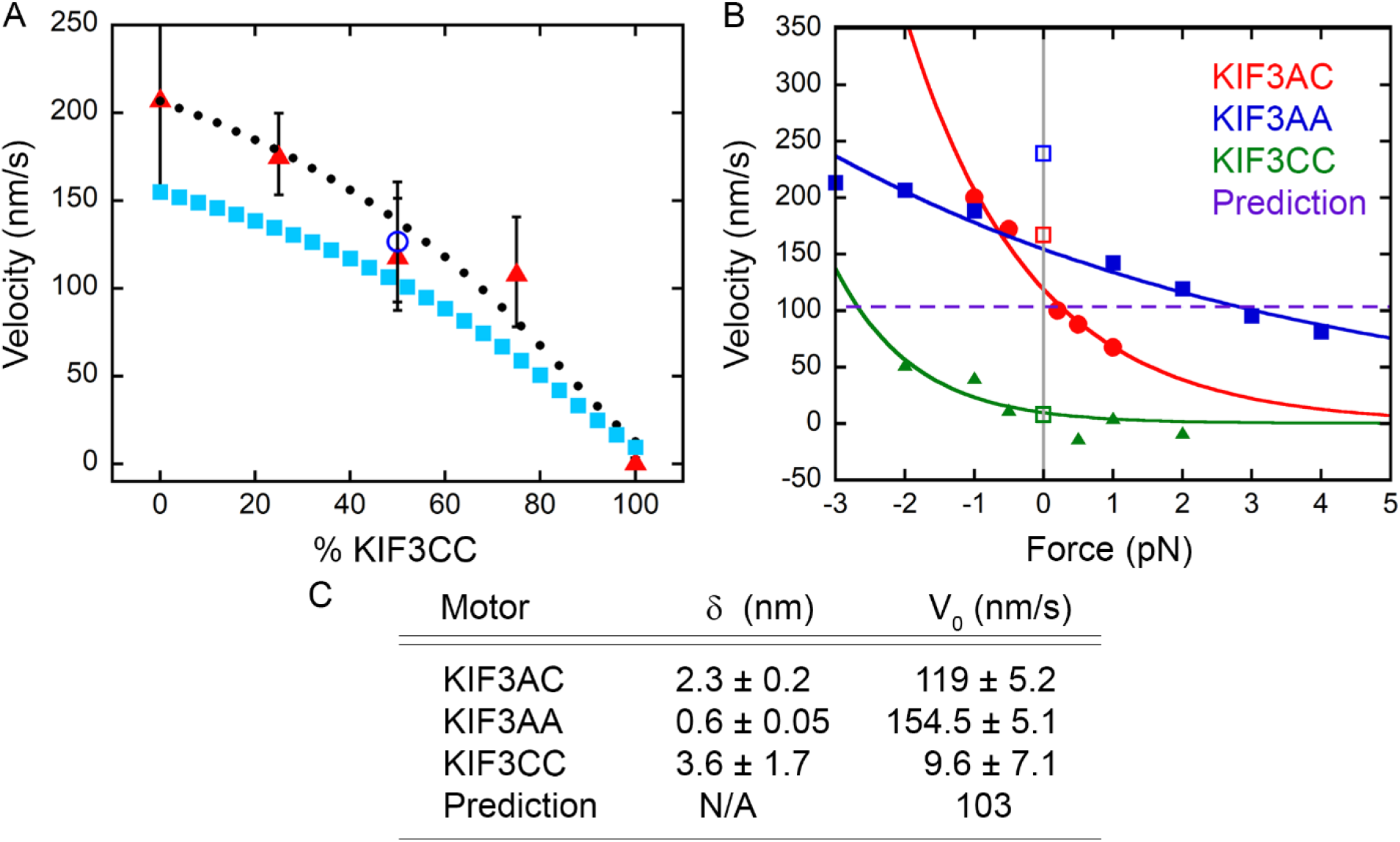
The unloaded velocity of KIF3AC is predicted by the force dependence of KIF3AA and KIF3CC velocities and recapitulated by MT gliding by mixtures of KIF3AA and KIF3CC. (A) Plot of MT velocity in a gliding filament assay versus percentage of KIF3CC observed for mixtures of KIF3AA and KIF3CC (red triangles) ± S.D. The gliding velocity observed for KIF3AC alone is shown (unfilled blue circle) ± S.D. The solid cyan squares represent the hypothetical predicted microtubule gliding velocity of any mixture of KIF3AA and KIF3CC based on the measured force dependence of the KIF3AA and KIF3CC homodimers in single-bead optical trapping assays, and the solid black circles show the predicted line multiplied by 1.3 to better align with the data. (B) Plot of velocity versus force with overlaid Bell equation fits for KIF3AC (red), KIF3AA (blue), and KIF3CC (green). The horizontal line (purple, dashed) crosses the KIF3AA curve at the same positive force as it crosses the KIF3CC curve in the negative force regime at ± 2.8 pN, and it intersects with the KIF3AC curve at zero force. (C) Parameters derived from the fit of the force dependence of velocity data for KIF3AC, KIF3AA, and KIF3CC. The prediction V_0_ represents the hypothetical velocity of KIF3AA and KIF3CC in a tug of war.

## Discussion

### KIF3AC steps at a rate that is distinct from KIF3AA or KIF3CC

How fast would KIFAC be expected to move if the two heads operated with the kinetics of their respective homodimers? With velocities of 270 and 7.3 nm/s for KIFAA and KIFCC, respectfully, the average dwell times between 8 nm steps are (*d* / *v* =) 0.029 s and 1.1 s. The KIFAC velocity predicted by the simplest scheme of half slow and half fast steps would be v = 2*(d nm) / (0.029 + 1.1 s) = 15 nm/s. At 110 nm/s, KIFAC clearly processively steps much faster than this predicted rate. Thus, the unloaded kinetic properties of KIF3AC are the result of heterodimerization, which impacts the properties of the motor domains in a way that is different from homodimerization.

A key finding of this study is that a dominant stepping rate (11 s^−1^) was observed for KIF3AC at 1 mM MgATP that is distinct from the stepping of KIF3AA (> 30 s^−1^) and KIF3CC (< 1.5 s^−1^) under identical conditions. The distribution of KIF3AC step durations is not a linear combination of fast KIF3AA and slow KIF3CC step durations, which would be clearly resolved in our experiments (Fig. *2A*). Previous work in which dimers were constructed with non-identical motors showed the alternating stepping rates expected from the alternating head kinetics (36). Even homodimeric kinesins have been reported to limp with alternating long and short dwell times before each step (37, 38). Thus, it was surprising that we did not observe alternating dwell times with KIF3AC. The high acceleration of the KIF3C head and only moderate deceleration of the KIF3A head in the KIF3AC dimer suggests kinetic tuning of the two heads, apparently driven by the strain-dependence of velocity of the individual heads.

### KIF3C can be activated by an assisting load or KIF3A

Although KIF3CC is an exceptionally slow motor in the absence of external force with little net plus-end movement under hindering loads, its stepping rate increases substantially with assisting loads (Fig. 6*B*). Notably, microtubule gliding in the presence of KIFCC is activated by the faster motor, KIF3AA (Fig. 7A), such that equal densities of KIF3CC and KIF3AA power gliding at a rate similar to KIF3AC. This result is similar to the intermediate speed observed with half-and-half mixtures of two *C. elegans* intraflagellar transport (IFT) *kinesins* with 3-fold different velocities (39). In contrast, gliding filament assays with mixtures of various combinations of myosin-II molecules with different velocities tended to be dominated by slower myosins (40, 41).

The shape of the microtubule gliding velocity curve as a function of the percent of KIF3CC is well modeled by assuming that forces proportional to the respective motor densities are assisting and hindering KIFCC and KIF3AA, respectively, and the kinetics of the motors are affected with the force-dependent parameters determined by optical trapping (Fig. 7*A*; see Methods). The rates predicted from the trap data are slightly slower than experimentally observed in gliding assays, which may be due to geometric differences between the two types of study (42). Notably, based on the force-dependent fits from the optical trapping experiments, an assisting load of −2.8 pN on KIF3CC is predicted to have the same speed as unloaded KIFAC (103 nm/s), which is the same speed as KIF3AA under 2.8 pN hindering load (Fig. 7*B*). Thus, the major effect of heterodimerization is the acceleration of KIF3C and slowing of KIF3A resulting in the observed speed. These results suggest the properties of KIF3AC are not likely due to “emergent” properties of the individual motor domains upon dimerization, but are due to differential mechanical constraints on the heads in the homo-and hetero-dimers.

### Intramolecular strain in KIF3CC is insufficient to activate efficient stepping

Intramolecular strain is a key component of motor-motor communication which enables high processivity and rapid stepping in kinesin dimers (43–47). Thus, we propose that intramolecular strain in the KIF3CC homodimer is insufficient to fully activate the stepping rate, and inclusion of KIF3A in the dimer generates the strain necessary to activate KIF3C to step more quickly. Indeed, the observed stepping kinetics of KIF3CC relative to KIF3AC mirror previous work showing that lengthening the neck-linker of kinesin-1 reduced the stepping velocity, yet assisting load could recover the velocity to the wild type value (43). Higher interhead compliance also increased the probability of backstepping of kinesin-1 (43), which is observed with our motor dimers that contain KIF3C (Figs. 2*C*, *6E and F*, Tables 1 and 4).

Intramolecular strain is transmitted via the neck-linkers in kinesin motors, yet the neck-linker domains of KIF3A and KIF3C are the same length (48). However, differences in the interactions between the neck-linker and the motor domains, possibly via differences in their cover-strand sequences (49, 50), result in stronger or weaker head-head coupling and ultimately affect ATPase kinetics (43–45, 51). When stepping against a stationary trap, KIF3CC detached at the lowest average force, KIF3AC detached at a higher average force while KIF3AA maintained the highest average force of the three. A previous study showed that decreasing intramolecular strain also caused kinesin-1 to detach or stall at lower forces than wild type (43). This observation supports the argument that low interhead strain drives the relatively low detachment force and velocity of KIF3CC. Reduced intramolecular strain may result in ungated detachment of the rearward head from the two-head-bound state. Such a motor would then be highly prone to detachment or stalling under hindering load.

### KIF3AC is an unconventional cargo transporter

It has been reported that transport of small axonal vesicles by kinesin-1 requires 2-3 motors (52, 53), yet transport by KIF3AC would likely require many more due to its relatively low force generating capabilities. A recent study suggested that scaling of total force generation with the number of motors may happen more efficiently for kinesin-2 motors than in other kinesin subfamilies, suggesting that these motors may be adapted to drive transport in larger teams (54). The high probability of KIF3AC detachment from the microtubule under load along with the fast microtubule association kinetics of KIF3AC (4, 17, 18) would enable KIF3AC to navigate obstacles on the microtubule effectively (55) or transport cargo more efficiently in larger ensembles (56), (57, 58)). Alternatively, the high probability of KIF3AC detachment from the microtubule under load may be an adaptation for cooperative transport with other faster motors as observed for KIF3AB-KAP and KIF17 (59). For these reasons, we argue that KIF3AC would be able to transport cargoes in a manner that is qualitatively different than other processive kinesins, and this difference is likely an adaptation for the specific biological role of KIF3AC.

## Methods

KIF3 motor constructs were expressed and purified as published previously (14). iSCAT, microtubule gliding (14), and optical trap motility assays were performed using standard motility assay conditions (see *Supplemental Appendix* **SI Methods** for details regarding assay conditions, instrumentation, and data collection). Kerssemakers’ algorithm was used to identify and fit steps in the data traces (31). Dwell time data were fit using maximum likelihood estimation in MEMLET (32) either to a single exponential function:

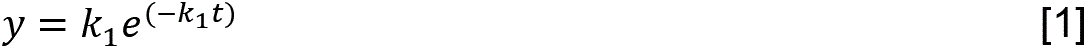

or to the sum of two exponential functions:

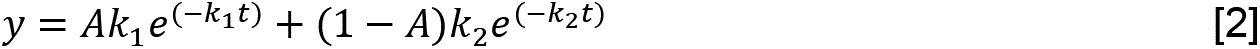

where A is the relative amplitude of the first component and (1 − A) is reported in the text as A_2_ but is not an independently fit variable. The dead time function in MEMLET was used to account for events that may not have been observed due to instrumental limitations. The hypothetical KIF3AA + KIF3CC dwell time cumulative distribution function (Fig. 2*A*) was calculated by taking 0.5 times the fit to the KIF3AA dwell time cumulative distribution function plus 0.5 times the fit to the KIF3CC dwell time cumulative distribution function. Step size data were fit using maximum likelihood estimation in the MEMLET software (32). Step size distributions derived from iSCAT experiments were fit as follows: Forward and backward steps were fit independently. A single Gaussian distribution was fit to the backward steps, and the sum of two Gaussian distributions to the forward steps, as justified by the log-likelihood ratio test. Step size distributions from optical trapping assays were fit by the sum of two Gaussian distributions. Step size data are reported as the mean ± σ of the fit curve, and σ describes the width of the Gaussian peak and not uncertainty in the peak position. Normalized step size histograms were plotted as probability density, and dwell time data were plotted as cumulative distributions (see **SI Methods** for detail). The Bell equation was fit to velocity versus force data (35):

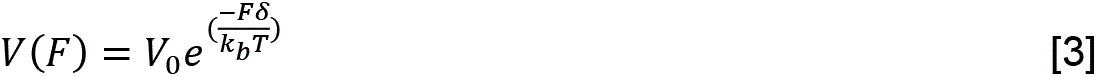

with *T* based on the assay temperature, 20 ± 1 °C, *V*_0_ is the unloaded velocity based on the fit, and the effective distance parameter *δ* defines the force sensitivity. In order for this form of the Bell equation to be valid, a constant average step size is assumed. The StatPlus plugin for Microsoft Excel (AnalystSoft Inc.) was used for statistical analysis for detachment, maximum, and comparison of stall forces was performed using an unpaired student’s *t* test with an α-reliability level of 0.05, and statistical analysis of stall percentage was performed using the “N−1” Chi-squared test with an α-reliability level of 0.05 (Table 2).

## Acknowledgements

We would like to express our gratitude to Dr. Henry Shuman for experimental advice, Dr. Philipp Kukura for providing advice and the software for operation of the iSCAT instrument, Dr. Matthew Caporizzo for training and input on iSCAT assays and data analysis, Dr. Scott Forth for valuable discussions, as well as current and past members of the Ostap, Goldman, and Gilbert labs for valuable discussion. This work was funded by NIH R37 GM054141 to S.P.G., NIH R35 GM118139 to Y.E.G. and NIH P01 GM087253 and Center for Engineering MechanoBiology NSF Science and Technology Center, CMMI: 15-48571 to Y.E.G. and E.M.O.

## Author Contributions

B.M.B., S.P., Y.E.G, S.P.G. and E.M.O designed the research; B.M.B. performed the experiments; B.M.B. and M.S.W. analyzed the data; B.M.B., M.S.W., S.P. and Y.E.G. wrote custom data analysis scripts; M.S.W. performed mathematical modeling; B.M.B., Y.E.G., S.P.G. and E.M.O wrote the paper; all of the authors read the paper and provided additional input.

## Declaration of Interests

The authors declare no competing financial interests.

