## Supplementary Materials for "Strain-Dependent Kinetic Properties of KIF3A and KIF3C Tune the Mechanochemistry of the KIF3AC Heterodimer"

**This PDF file includes:**

Supplementary Results
Supplementary Methods
Figures S1 to S5
SI References

### Supplementary Results

**Mathematical models impose limits on KIF3AC kinetics.** Based on the single molecule velocities reported here and previously (1, 2), we propose that the KIF3C head has become faster in KIF3AC because for a motor stepping with alternating rates, the slowest that the slow motor can step is approximately one half the observed velocity of the heterodimer. Therefore, at the observed KIF3AC stepping velocity at 1 pN hindering load of ~70 nm/s (~9 $s^{-1}$) for KIF3AC, the slowest that that KIF3C could step in this context would be ~4.5 $s^{-1}$ based on velocity. Limits can also be placed on the possible stepping rate constants by simulations of dwell times to determine under which conditions a single exponential distribution would be observed even if the distribution was composed of dwell times with two independent rates. Simulations were performed in which an equal combination of dwell times were generated at two distinct rates, with one rate fixed at 8 $s^{-1}$ and the other varied from 1 to 20 $s^{-1}$. In these simulations a double exponential function was only statistically justified by the log-likelihood ratio test when the difference in the two rates was >2-fold (*Supplementary Appendix* Fig. S4*A* and *B*). Additionally, an equal combination of dwell times at 8 $s^{-1}$ and 15 $s^{-1}$ was simulated with the same number of points as the real dataset for KIF3AC stepping against a 1-pN hindering load. This simulated dwell time distribution was best fit by a single exponential function according to the log-likelihood ratio test (*Supplementary Appendix* Fig. S4*C*), which shows that a measured single exponential rate could be the result of two rates in combination, provided that the fold difference between these rates is <2. This requirement places upper and lower bounds on the possible stepping rates in KIF3AC under these conditions, ranging from both steps occurring at ~11 $s^{-1}$, or two distinct rates with slower rate between 8 and 11 $s^{-1}$ and the faster rate between 11 and 15 $s^{-1}$. These results show that, for KIF3AC, steps by KIF3C became faster, and steps by KIF3A became slower than in the homodimers.

**Supplementary Methods**

**KIF3 Motor Constructs.** The *M. musculus kif3a, kif3b,* and *kif3c* plasmids for expression of KIF3AB and KIF3AC heterodimers as well as the engineered KIF3AA and KIF3CC homodimers were described previously (1). Each motor construct contains the native sequence of the motor domain, neck linker, and helix $\alpha$7 (KIF3A, M1-L374; KIF3B, M1-K371; KIF3C, M1-L396) followed by either the EB1 dimerization motif or an acidic/basic heterodimerization domain (AHD/BHD) in bold (below), the Tobacco Etch Virus (TEV) protease-cleavable site (italics), a linker (plain font), and either the StrepII or $His_8$ tag (underlined). The amino acid sequence of KIF3A used to generate KIF3AC heterodimers is as follows:

KIF3A(M1-L374)-

**DFYFGKLRNIELICQENEGENDPVLQRIVDILYATDE**TTS*SENLYFQ*GAS<u>NWSHPQFEK</u>.

To generate homodimers of KIF3AA, the C-terminal StrepII tag of this construct was exchanged for a $His_8$ tag. The amino acid sequence of KIF3C used to generate KIF3AC heterodimers is as follows:

KIF3C(M1-L396)-

**DFYFGKLRNIELICQENEGENDPVLQRIVDILYATDE**TTS*SENLYFQ*GAS<u>HHHHHHHH</u>.

To generate homodimers of KIF3CC, the same construct was used. To generate the KIF3AC$\Delta$L11 heterodimers, the KIF3C construct had the following modifications: P258A, $\Delta$N259-S284. The EB1 motif used to generate KIF3AC heterodimers and KIF3AA and KIF3CC homodimers is a dimerization domain only and does not have MT binding activity (1). The TEV-cleavable purification tags remained intact for all experiments.

For KIF3AB, a synthetic heterodimerization motif (SHD) containing either an acidic heterodimerization domain (AHD) or basic heterodimerization domain (BHD) was used. The amino acid sequence of KIF3A used to generate KIF3AB heterodimers is as follows:

KIF3A(M1-L374)- **LEKEIAALEKEIAALEK**TTS*ENLYFQ*GASNWSHPQFEK.

The amino acid sequence of KIF3B used to generate KIF3AB heterodimers is as follows: KIF3B(M1-K371)- **LKEKIAALKEKIAALKE**TTS*ENLYFQ*GASHHHHHHHH.

The KIF3A construct contains the AHD sequence and the KIF3B construct contains the BHD sequence. Previous experiments showed that there was no difference in the single molecule results for heterodimers stabilized by the SHD versus EB1 (1).

**KIF3 Motor Expression and Purification.** All KIF3 motors were expressed in *E. coli* BL21-CodonPlus (DE3)-RIL cells (Strategene). Heterodimers resulted from cotransformation of two plasmids, each with different antibiotic resistance, and transformants were plated on triple-selective LB plates containing 100 μg/mL ampicillin, 50 μg/mL kanamycin, and 10 μg/mL chloramphenicol. To purify exclusively heterodimeric KIF3AB or KIF3AC, sequential affinity columns were used. The HisTrap FF $Ni^{+2}$-NTA column (GE Healthcare) was used to select for the C-terminal $His_8$ tags of KIF3B or KIF3C, followed by the StrepTactin column (GE Healthcare), which selected the C-terminal StrepII tag of KIF3A. Purified KIF3AB or KIF3AC motors were assessed by analytical gel filtration chromatography (Superose 10/300; GE Healthcare) and SDS-PAGE to confirm a single population of heterodimeric motors with a 1:1 stoichiometry of KIF3A to KIF3B or KIF3C. The predicted molecular weight of KIF3AB heterodimers is 92,131, and the predicted dimer molecular weight of KIF3AC is 98,317 based on amino acid sequence.

To purify homodimeric KIF3AA or KIF3CC, the supernatant of a cell lysate expressing KIF3A or KIF3C was loaded onto the HisTrap FF $Ni^{+2}$-NTA column (GE Healthcare) to select for $His_8$-tagged protein. Fractions with KIF3AA or KIF3CC were pooled and additionally purified by size exclusion chromatography (Superose 10/300 column; GE Healthcare) and eluted into dialysis buffer (20 mM HEPES pH = 7.2 with KOH, 200 mM NaCl, 5 mM magnesium acetate, 0.1 mM EDTA, 0.1 mM EGTA, 50 mM potassium acetate, and 1 mM DTT plus 5% w/v sucrose). The purity of homodimeric KIF3AA and KIF3CC was assayed using analytical gel filtration and SDS-PAGE. KIF3AA has a predicted dimer molecular weight based on sequence of 97,000, and KIF3CC has a predicted dimer molecular weight of 99,518.

Purified KIF3 dimers were dialyzed against ATPase buffer (20 mM HEPES pH = 7.2 with KOH, 5 mM magnesium acetate, 0.1 mM EDTA, 0.1 mM EGTA, 50 mM potassium acetate, and 1 mM DTT plus 5% w/v sucrose), clarified by ultracentrifugation to remove aggregated protein, and aliquoted at experimental volumes. Aliquots were flash-frozen in liquid nitrogen and stored at -80 °C. Before each experiment, KIF3 protein aliquots were thawed rapidly and clarified for 10 min at 4 °C and 313,000 × *g* (TLA-100 rotor, Beckman Coulter TLX Optima Ultracentrifuge). Protein concentrations were determined using the Bio-Rad protein assay with IgG as the protein standard.

**Rigor Kinesin Construct and Purification.** A rigor mutant of human KIF5B encoding the first 560 amino acid residues of KIF5B with the point mutation G234A, the gift of Dr. Linda Wordeman, was used to adhere microtubules to the coverslip surface in iSCAT motility assays (3). *E. coli* (BL21) were used for expression of rigor kinesin dimers. The supernatant of a cell lysate expressing KIF5B G234A was loaded onto a 3-ml $Co^{2+}$ resin column (TALON) and eluted with elution buffer (300 mM NaCl, 250 mM Imidazole HCl pH = 7.0, 1 mM $MgCl_2$, 10 μM MgATP with 10% v/v glycerol). Protein aliquots were flash-frozen in liquid nitrogen and stored at -80 °C with protein concentration determined by Bradford Assay.

**Microtubule Preparation.** Unlabeled porcine brain tubulin (Cytoskeleton, Inc.) was diluted to 5 mg/ml in BRB-80 pH = 6.8 (80 mM PIPES pH = 6.8, 1 mM EGTA, 1 mM $MgCl_2$) plus 1 mM GTP on ice. Rhodamine-labelled tubulin (Cytoskeleton, Inc.) was reconstituted in BRB-80 pH = 6.8 plus 1 mM GTP on ice at a concentration of 5 mg/ml. Twenty-four parts unlabeled tubulin were mixed with one part rhodamine-labelled tubulin and incubated on ice for 5 min prior to clarification by ultracentrifugation for 10 min at 4 °C at 313,000 × g (TLA-100 rotor, Beckman Coulter TLX Optima Ultracentrifuge). The resulting supernatant was incubated at 37 °C for 20 min followed by supplementation with 75 μM final concentration paclitaxel (Cytoskeleton, Inc.) and incubation for an additional 10 min at 37 ºC. The microtubules were subsequently adjusted to 110 μM paclitaxel followed by a final incubation for 10 min at 37 °C. Assembled microtubules were pelleted for 20 min at 25 °C at maximum speed in a tabletop centrifuge. The microtubule pellet was resuspended in BRB-80 pH = 7.5 plus 40 μM paclitaxel at a final concentration of ~41 μM tubulin dimer. Polarity-marked microtubules for microtubule gliding assays were prepared as described previously (4).

**Microtubule Gliding Assays.** Microtubule gliding assays were performed in perfusion chambers made from an acid-washed coverslip mounted on a glass slide separated by strips of double-sided tape to generate a flow cell of ~10-μl volume. Anti-His antibody was diluted into BRB-80 at a final concentration of 100 μg/ml and flowed into the perfusion chamber for 5 minutes. Next, chambers were blocked with BRB-80 plus 1 mg/ml casein, 0.2 mg/ml glucose oxidase, 0.175 mg/ml catalase, 25 mM glucose, and 20 μM paclitaxel. MT•motor complex was preformed (500 nM polarity-marked MTs, 2.5 μM total motor, and 20 μM paclitaxel) and flowed into the chamber for 10 minutes. The chamber was then washed with blocking buffer, and final buffer (BRB-80 plus 1 mM MgATP, 0.3 mg/ml creatine phosphokinase, 2 mM phosphocreatine, 1 mg/ml casein, 0.2 mg/ml glucose oxidase, 0.175 mg/ml catalase, 25 mM glucose, and 20 μM paclitaxel) was flowed into the chamber prior to sealing the chamber with clear nail polish. MT gliding assays were imaged using a Zeiss Inverted Axio Observer Z1 MOT fluorescence microscope with the 100X oil 1.46 N.A. Plan-Apochromat objective (Carl Zeiss Microscopy, Jena, Germany). Images were collected via a Hamamatsu EM-CCD camera driven by the Zen software package. 512 x 512 pixel images were captured every 20 s for 20 minutes (KIF3AA, KIF3AC, and mixtures) or every 30 s for 30 minutes (KIF3CC). All experiments were performed with same final motor concentration of 2.5 μM motor dimer. Gliding microtubules were tracked with the MTrackJ plugin for ImageJ (5). To be scored as a gliding microtubule, a microtubule must (a) be between 3 and 8 μm in length, (b) remain in the field of view and maintain a constant velocity for at least 5 frames, and (c) not be observed to stall or detach from the surface during the time course of imaging.

**iSCAT Motility Assays.** Streptavidin-coated 50-nm gold nanoparticles (Nanopartz, Inc.) were incubated with an excess of biotinylated anti-His antibody (Qiagen) on ice for 1 hour. A 3 nM working concentration of nanoparticle-antibody complex was generated from the stock nanoparticles (3.3 nM) and anti-His antibody (200 μg/ml). Nanoparticle-antibody complex was then incubated with 300 pM kinesin motor on ice for 15 min. Final concentrations: 2.4 nM nanoparticle-antibody complex, 240 pM kinesin dimer. Motility assay perfusion chambers were prepared with a plasma-cleaned 22x40 mm coverglass adhered to a glass slide with double-sided sticky tape and a typical flow cell volume of ~20 μl. Rigor kinesin diluted to 0.1-0.3 mg/ml in BRB-80 pH = 7.5 was flowed into the perfusion chamber and incubated for 5 min prior to two washes with BRB-80 plus 1 mg/ml casein. Microtubules were diluted 1:300 (~136 nM final concentration) in BRB-80 plus 20 μM paclitaxel, flowed into the perfusion chamber and incubated for 10 min prior to a wash with BRB-80, pH = 7.5 plus 20 μM paclitaxel, 20 mM DTT, 240 pM gold nanoparticles with 24 pM KIF3 dimer, and incubated for 5 min. The final motility mix (BRB-80, pH = 7.5, 20 μM paclitaxel, 0.1 mg/ml catalase, 0.2 mg/ml glucose oxidase, 20 mM glucose, 20 mM DTT, 1 mM $MgCl_2$, 1 mM ATP, and 1 mg/ml casein) was flowed in immediately prior to imaging. Perfusion chambers were imaged for a maximum of 5 min to minimize photodamage to the sample.

The iSCAT instrument was custom built and driven by custom LabVIEW software (National Instruments). Data were acquired at 1,000 frames per second and 10 seconds per video. The iSCAT instrument utilized an OBIS 488-nm LS FP diode laser (Coherent), MV1-D1024E-160-CL-12 CMOS camera (Photon Focus), and 1.49 NA CFI Apochromat TIRF 100X oil immersion objective (Nikon) (6-10). This imaging strategy yielded 256x256 pixel images with a pixel size of 83.4 nm/pixel. The laser intensity was adjusted until the reflection from the slide approximately half-saturated the camera digitizer at the working frame rate.

**iSCAT Image Analysis and Particle Tracking.** Images were initially analyzed in the FIJI software package (11), where regions of interest (ROIs) containing motility events were identified, cropped, background subtracted, and saved. ROIs were imported into the FIESTA program (12) in MATLAB® (Mathworks) for particle tracking. Intensity profiles of tracked particles were fit to a 2D

Gaussian distribution, with user input initial guesses of a 5- to 10-pixel particle area and full width at half maximum of 300 nm. Variable particle intensity and no filtering for thresholding were used for particle identification. A second order polynomial was fit to the particle trajectory to determine the path of travel, and the distance travelled by the particle along the path was used for further analysis. iSCAT motility velocities were determined from a linear fit to the distance along path of travel versus time. For step-fitting analysis, iSCAT positions along path of travel data were filtered with the Chung-Kennedy filter (13) with a total filter width of 11 ms, defined as data 5 ms ahead of and 5 ms behind the central point, and a sensitivity factor of 2.

**Optical Trap Motility Assays.** Streptavidin-coated 0.82 μm polystyrene beads at a stock concentration of $3.55 \times 10^{10}$ beads per ml (Spherotech) were used. To prepare beads for motility assays, beads were diluted 5-fold in BRB-80, pH = 7.5, bath sonicated for 5 min and diluted 2-fold further to a final concentration of $3.55 \times 10^{9}$ beads per ml in BRB-80. Biotinylated anti-His antibody (Qiagen) was added to a final concentration of 1 μg/ml and the reaction was incubated overnight at 4 ºC with constant rotation. Beads were washed 3 times by pelleting at low speed and resuspension in BRB-80 plus 1 mg/ml casein. Prior to the optical trap motility assays, beads were incubated with KIF3 motor for 15 min at 4 °C. Appropriate motor concentration for single molecule conditions was determined by titration such that ~1 out of every 10 beads produced an interaction with the microtubule, with typical motor concentrations in the incubation step of 50 to 500 pM kinesin dimer.

Nitrocellulose-coated chambers were prepared with 0.1% nitrocellulose solution (EMS) and allowed to air dry. Coverslips were adhered to each other with double sided tape to form ~20 μL perfusion chambers. 175 μg/mL mouse anti-tubulin antibody (BioRAD) in BRB-80 pH = 7.5 was flowed into the chamber and incubated for 5 min, followed by blocking with 2 mg/ml casein in BRB-80 for 5 minutes. ~100 nM rhodamine-labelled microtubules in BRB-80 with 20 μM paclitaxel and 1 mg/ml casein was flowed into the chamber and incubated for 5 min prior to a subsequent wash with BRB-80 plus 20 μM paclitaxel and 1 mg/ml casein. Bead:motor complex was diluted 1:25 in the final motility buffer (BRB-80 with 20 mM glucose, 0.1 mg/ml catalase, 0.2 mg/ml glucose oxidase, 20 μM paclitaxel, 20 mM DTT, 1 mM $MgCl_2$, 1 mM ATP and 1 mg/ml casein) and flowed into the chamber prior to sealing the chamber with vacuum grease. The final chamber concentration of kinesin dimer was between 2 and 20 pM.

Ramp-force measurements were executed on the optical trap instrument as described (14, 15) but with a 63X magnification, 1.2 N.A. water immersion objective (Zeiss). Experiments were performed at room temperature (20 ± 1 °C). The trap stiffness was calibrated based on the power spectrum of the thermal motion of a trapped bead, and calibrations were done for each bead used for data collection. Data collection was accomplished through custom LabVIEW software (14, 15) (National Instruments).

Force-feedback experiments were performed on the optical trap instrument described previously (16, 17), except that one of the two laser beams was blocked. A 1064-nm trapping laser was used with a trap stiffness of approximately 0.05 pN $nm^{-1}$, determined from the power spectrum of the thermal motion of a trapped bead. Force-feedback was conducted using a digital feedback loop and an electro-optical deflector (EOD) to steer beam position and maintain constant force on the trapped bead during a kinesin-driven motility event (16). Hindering loads were generated by the beam following the movement of the trapped bead, and assisting loads were generated by the beam leading the movement of the trapped bead. Data were acquired at 200 kHz, and the digital feedback loop calculations were performed at 200 kHz. Data acquisition, feedback calculations, and beam position control output were conducted by a LabVIEW Multi-function I/O device with built-in FPGA (PXI-7851) and custom written LabVIEW software (16).

**Optical Trap Data Analysis.** Ramp force data acquired from the optical trap were analyzed with custom LabView code. The custom script identified ramps, measured the detachment force and maximum force during the ramp, and calculated the mean force and standard deviation of force over the final 70 ms of the ramp. This mean was used to define the stall force as long as the standard deviation of the force over that window was within 5% of the mean force. If the standard deviation of the force was greater than 5% of the mean, the event was considered not to end in a stall. Detachment force was defined as force on the bead at the moment of detachment from the microtubule, and maximum force was defined as the greatest magnitude force reached during the event. Definitions of detachment, maximum, and stall force were derived from (18). The mean detachment, maximum, and stall forces were plotted as histograms. The stall percentage is defined as the number of events that terminated in a stall divided by the total number of events identified, expressed as a percentage.

Constant force position versus time ramp data was analyzed using custom MATLAB® software (Mathworks). Both raw and filtered data were plotted versus time to aid in event identification. Raw data were acquired at 200 kHz, then were down-sampled by averaging with a 10-point window prior to filtering and plotting. Data were filtered using the Chung-Kennedy (13) filter with a total width of 3.05 ms, meaning the filter used 1.5 ms forward and 1.5 ms behind the central point and a sensitivity factor of 2. Ramps were identified carefully by eye, based on the following criteria: (1) the plotted force value was stable at the force setpoint and (2) the excursion from the initial position was greater than 16 nm. To avoid inclusion of events where the motor slipped toward the minus end of the microtubule and reattached without returning to the initial trap position, any rearward motion of ≥16 nm was considered the end of an event. A new event began at the point of reattachment. Deviation in the force value from the setpoint was also indicative of detachment and marked the end of a ramp. Run lengths of events were determined based on the position of the final data point minus the position of the first data point, and velocities were determined based on the linear fit to the trajectory. For any given condition, 100-250 events were identified and used to quantify run length and velocity.

**Step Size and Kinetics Analysis.** The Kerssemakers' algorithm was used to identify and fit steps in position versus time traces from iSCAT and optical trap motility assays (19). The algorithm was run in MATLAB®. Steps were fit to filtered data. In order to minimize under- or over-fitting, each event was refit with 10% more of 10% fewer steps to show that mean step size and dwell time were minimally sensitive to the number of identified steps.

**Data Fitting.** Velocity, run length, step size, and step dwell time data were fit using MEMLET software (20). A Gaussian distribution was fit to velocity data in order to determine their means and standard deviation. A single exponential decay probability density function was fit to run length data to determine the mean run length:

$$y = \frac{1}{r} e^{\frac{-L}{r}} \quad [1]$$

where the mean run length is $r$. The standard deviation of run length populations were calculated with Microsoft Excel. A 16-nm minimum threshold run length, the shortest event that could be positively identified, was used for determining mean run length through fitting with MEMLET.

Step size data were plotted as a normalized histogram with the y-axis as probability density. This plotting method yields bins where the area of any given bin is defined as the probability that the step size lies between the upper and lower bounds of that bin;

$$Bin\ Area = P(x_1 < X < x_2) \quad [2]$$

where $X$ is the randomly distributed variable being plotted (step size), $x_1$ is the bin lower limit, and $x_2$ is the bin upper limit. The height of a bin (probability density) is equal to the bin area divided by the bin width:

$$Probability\ Density = \frac{P(x_1 < X < x_2)}{(x_2 - x_1)} \quad [3]$$

The sum of the areas of the bins is 1 (20). The sum of two Gaussian distributions was fit to step size distributions derived from optical trap assays. Forward and backward steps observed by iSCAT microscopy were fit independently. Backward steps were fit to a Gaussian distribution, and forward steps were fit to the sum of two Gaussian components.

Either a single exponential decay function or the sum of two exponential functions was fit to dwell time data, with justification of a second exponential based on the log-likelihood model testing function in MEMLET. The sum of two independent exponential functions has the form:

$$y = Ak_1 e^{(-k_1 t)} + (1 - A)k_2 e^{(-k_2 t)} \quad [4]$$

where $k_1$ and $k_2$ reflect the rate constants of the two exponential decay functions. All probability density functions used here are built into the MEMLET software. Dwell time data were fit with a dead time of 20 ms, which is equal to twice the period of the smoothing filter applied to the data. Dwell time data were plotted as cumulative distribution function plots.

**Prediction of Gliding Velocity as a Function of KIF3CC Proportion in Homodimer Mixtures.** The expected velocity of microtubule gliding powered by mixtures of KIF3AA and KIF3CC was modeled based on the relative proportions of KIF3AA and KIF3CC as well as their force dependences. Since the velocity is constant, the net force on a gliding microtubule is zero, the drag force on KIFAA motors caused by the slower KIFCC motors being balanced by the assisting force on the KIFCC motors produced by the faster KIFAA motors. Inverting the Bell equation to give force as a function of velocity and totaling the two forces to zero gives $\alpha \frac{k_B T}{\delta_{CC}} \ln\left(\frac{V}{V_{0CC}}\right) + (1-\alpha)\frac{k_B T}{\delta_{AA}} \ln\left(\frac{V}{V_{0AA}}\right) = 0$, where $V$ is the predicted velocity for a mixture of ($\alpha$) KIFCC and (1-$\alpha$) KIFAA, and the $\delta$ and $V_0$ values for the two homodimers are given in Table of Fig. 7. Solving for V gives

$$V(\alpha) = \exp\left[\frac{\frac{\alpha \ln(V_{0CC})}{\delta_{CC}} + \frac{(1-\alpha)\ln(V_{0AA})}{\delta_{AA}}}{\frac{\alpha}{\delta_{CC}} + \frac{(1-\alpha)}{\delta_{AA}}}\right] \qquad [5]$$

which is plotted as the blue squares in Fig. 7A.

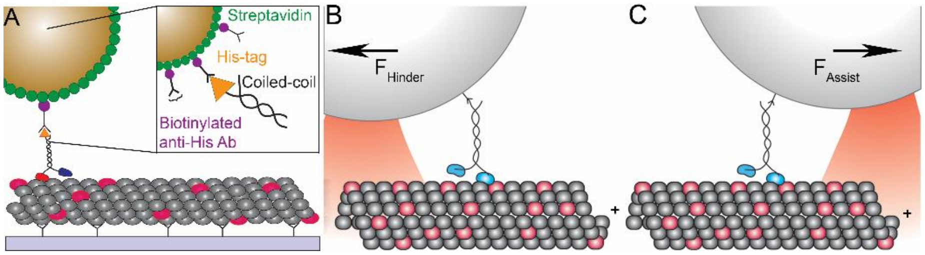


**Fig. S1. iSCAT attachment strategy and assisting and hindering constant load assay geometry.** (A) Related to Fig. 1. Illustration of attachment strategy used in iSCAT motility assays. Streptavidin-coated gold nanoparticles were incubated with a biotinylated anti-His antibody (Ab) which bound to the C-terminal $His_8$-tag of the KIF3 motors. (B and C) Related to Fig. 4. Illustration of the geometry of constant hindering and assisting load motility assays. Under constant hindering load (B) a digital feedback loop is used to maintain a constant displacement between the bead and trap center as the trap follows the walking motion of the kinesin attached to the bead, applying a force opposing the motion of the kinesin. With a constant assisting load (C) the feedback loop maintains a constant displacement between the bead and trap center as the trap leads the walking motion of the kinesin attached to the bead, applying a force assisting the motion of the kinesin.

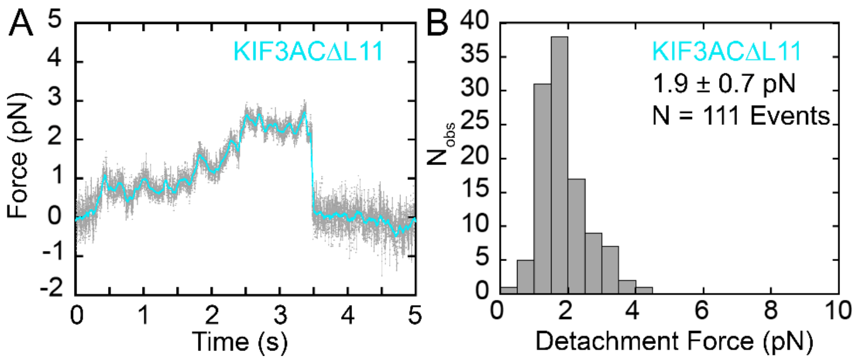


**Fig. S2. The extension of Loop L11 of KIF3C does not regulate force sensitivity in KIF3AC. Related to Fig. 3.** Sample ramp (A) of KIF3ACΔL11 stepping against a stationary trap. Force on the bead increases as the motor steps away from the trap center. Histogram of detachment force (B) observed for KIF3ACΔL11. Detachment force is defined as the amount of force on the bead immediately prior to detachment. $F_{detach}$ is displayed as mean ± S.D.

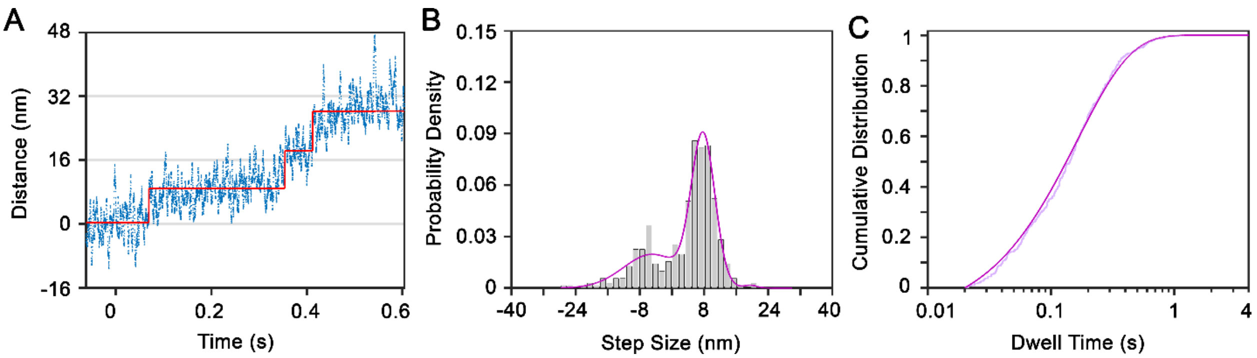


**Fig. S3: KIF3AC steps with a single rate under load in presence of rate-limiting MgATP.** Related to Fig. 6. (A) Sample trace of KIF3AC stepping against a 1-pN hindering load at 10 μM MgATP. (B) Histogram of step sizes observed for KIF3AC in the conditions listed for (A). (C) CDF plot of dwell times for KIF3AC in the conditions listed for (A). Lighter trace represents the data with the darker overlay as the single exponential fit to the data. $N_{steps}$ = 435.

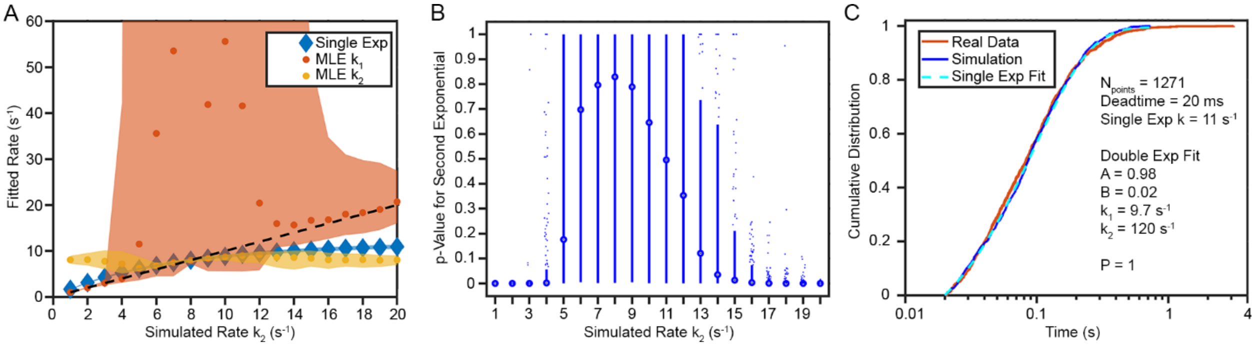


**Fig. S4.** Related to Fig. 2 and 6. (A) Simulations were carried out to address the question of whether a measured single exponential rate constant could result from a mixture of two similar but distinct rates. Dwell times from a two-exponential distribution were simulated with equal proportions of dwells from each of two rates, $k_1$ and $k_2$. Simulated dwell time distributions were fit by both a single exponential function and the sum of two exponential functions. One exponential rate $k_1$ was fixed at 8 $s^{-1}$ in all simulations while the second rate $k_2$ was varied from 1 to 20 $s^{-1}$. In each simulation 1200 dwell times were generated, and 200 independent simulations were completed to determine confidence intervals. Each data point represents the mean of the rates $k_1$ (red) and $k_2$ (goldenrod) derived from the fit of a two-exponential function to each simulated dataset. The mean rate from a single exponential fit is shown in blue. The 90% CI of each mean is shown as the shaded area and the dashed line represents the true simulated $k_2$ values. When the fold difference between $k_1$ and $k_2$ is small, $k_2$ is poorly constrained and the fit value diverges from the true simulated rate. (B) Box plot of p-values from the log-likelihood ratio test in MEMLET, indicating whether a second exponential component is statistically justified versus simulated rate $k_2$. When the difference between $k_1$ and $k_2$ is less than 2-fold, fitting a second exponential is often not observed to be statistically justified ($p>0.05$). (C) An even mixture of dwell times coming from 8 $s^{-1}$ and 15 $s^{-1}$ rates were simulated, giving the same number of points as in the real data set shown in red (KIF3AC with a 1-pN hindering load at 1 mM MgATP). Fitting these data shows that a single exponential function is a good fit to the data, and that fitting the sum of two exponential functions to the data is not statistically justified according to the log-likelihood ratio test. Fit parameters are shown in figure inset.

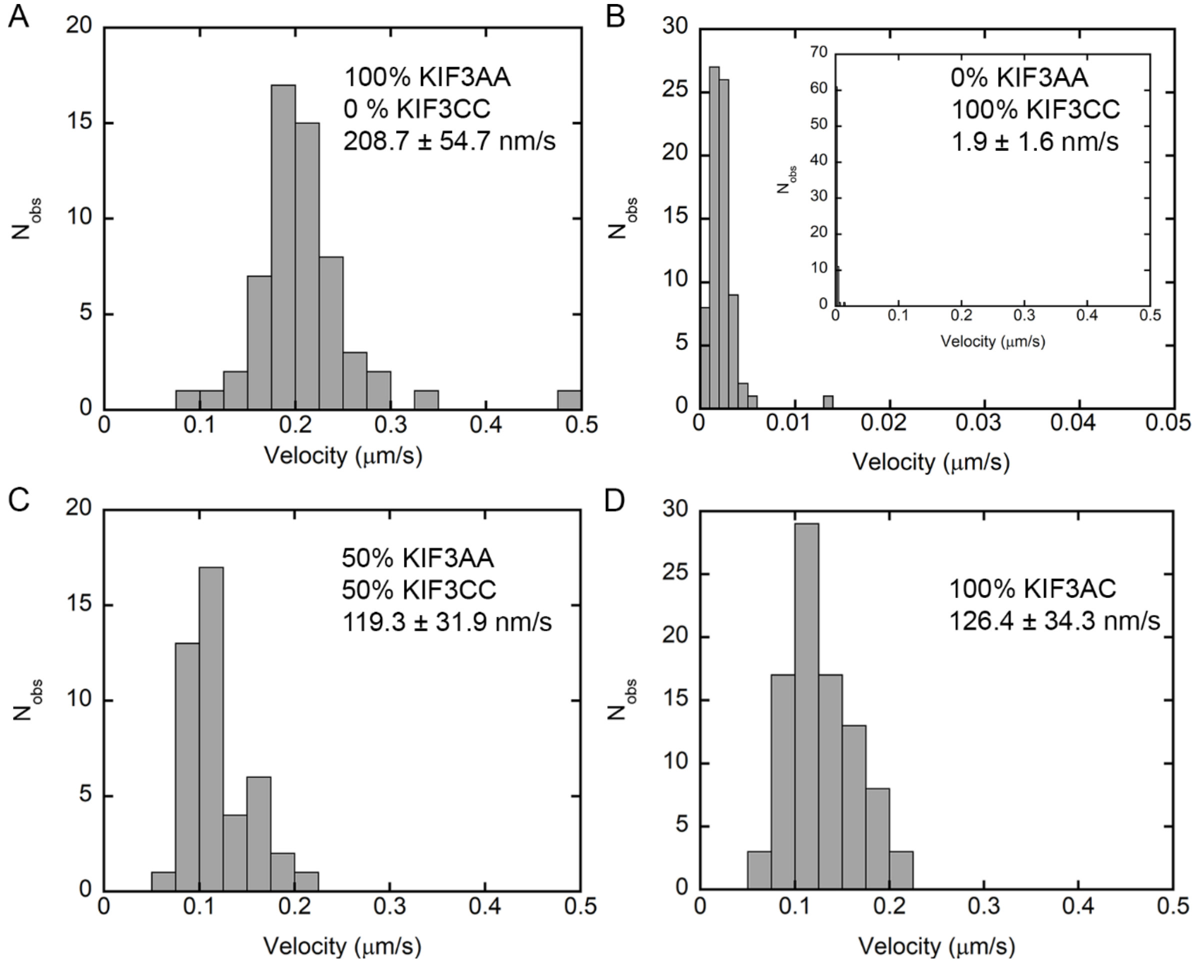


**Figure S5. Mixing KIF3AA and KIF3CC in MT gliding assays reduces gliding velocity in a manner dependent on relative KIF3CC concentration and can recapitulate the rate of KIF3AC MT gliding.** Related to Fig. 7*A*. (A-D) Histograms of MT gliding velocities observed for KIF3AA alone, N = 58 MTs (A), KIF3CC alone, N = 73 MTs (B), an equal mixture of KIF3AA and KIF3CC, N = 44 MTs (C), and KIF3AC alone, N = 90 MTs (D). KIF3CC data are shown on an expanded x-axis due to the ~30-fold difference in gliding velocity between KIF3CC and KIF3AA.
